# Cytokines driven by caspase-1 and RIPK3 are antagonistic

**DOI:** 10.64898/2026.06.19.733458

**Authors:** Lupeng Li, Jinyi (Sammy) Guo, Ying Wang, Carolyn A. Lacey, Yaxin Liu, Jianxiang Peng, Yu-Ming Li, Heather N. Larson, Melanie N. Pratt, Bart Tummers, Douglas R Green, Edward A. Miao

## Abstract

Pyroptosis, apoptosis, and necroptosis are thought to provide redundant protection against intracellular pathogens. We found that mice lacking inflammasomes, caspase-8, and MLKL were highly susceptible to infection by an environmental cytosol-invasive bacterium. Surprisingly, deletion of *Ripk3* completely restored resistance despite the continued absence of all three cell death pathways. The infection outcome was determined instead by the production of antagonistic cytokines arising from these pathways. Deletion of *Casp8* and *Mlkl* initiated an effector-triggered immunity-like response, in which RIPK3 induced type I interferons causing catastrophic susceptibility by impeding neutrophil responses. Opposing this, caspase-1-dependent IL-1β production counteracted these type I interferons and restored resistance. Thus, susceptibility arose not from failure of regulated cell death, but from the activation of RIPK3-driven type I interferons unopposed by caspase-1-driven IL-1β. Therefore, antagonistic cytokines embedded within cell death pathway signaling, distinct from cell death itself, can dictate the immune response to intracellular infection.

## Introduction

Pyroptosis, apoptosis, and necroptosis are regulated cell death pathways that collectively contribute to innate immune defense against intracellular pathogens^1–4^. These pathways eliminate intracellular replication niches while simultaneously generating inflammatory signals that recruit and activate additional immune effectors. Pyroptosis is initiated by inflammasomes, which are multiprotein complexes assembled by cytosolic pattern-recognition receptors (e.g., NLRP3 and AIM2). After these inflammasomes detect infection, they signal through the ASC adaptor protein to activate caspase-1, which cleaves and activates the pore-forming protein gasdermin D (GSDMD) to cause pyroptosis^5,6^. Extrinsic apoptosis is mediated by caspase-8, whereas necroptosis is driven by RIPK3-dependent phosphorylation of MLKL^7–9^. Extensive crosstalk among the pathways that drive these three forms of cell death suggests that they function as an integrated defense network with multiple avenues for redundancy^10^. Studies have demonstrated that these pathways can function redundantly during bacterial infection: simultaneous disruption of cell death signaling results in profound susceptibility to intracellular pathogens including *Salmonella* and *Shigella* species^11,12^. The prevailing model is that elimination of infected cellular niches, regardless of the mode of cell death, is the primary antimicrobial defense mechanism against intracellular infection^10,13^.

Regulated cell death pathways also generate multiple outputs beyond cellular destruction, including cytokine production, inflammatory signaling, and immune-cell recruitment. Caspase-1, for example, simultaneously drives pyroptotic cell death and the maturation and release of IL-1β and IL-18^5,14^. These outputs are mechanistically coupled yet serve distinct protective functions during infection. The relative contribution of cell death versus cytokine signaling during bacterial infection has not been fully studied.

Environmental opportunistic pathogens provide a powerful tool for addressing fundamental questions in innate immunity^4,15^. Unlike host-adapted pathogens that have evolved extensive mechanisms to evade mammalian immunity, environmental bacteria are rapidly eliminated by non-permissive hosts. As a result, the pathways responsible for their clearance are precisely those that constitute effective innate defense, making environmental bacteria particularly informative models. Studies of environmental bacteria have yielded fundamental insights into innate immunity: investigation of *Burkholderia thailandensis* led to the identification of caspase- 11 as a cytosolic lipopolysaccharide sensor^16,17^, while study of *Chromobacterium violaceum* provided compelling in vivo evidence for NLRC4 function^18^, uncovered a unique cytoprotective role for caspase-7 in membrane repair^19,20^, and established an innate granuloma model in which GSDMD is required for defense^21,22^. In each case, the absence of sophisticated pathogen immune evasion allowed clear visualization of otherwise obscured host defense mechanisms.

*Francisella philomiragia* is an environmental Gram-negative bacterium that can cause opportunistic infections in immunocompromised individuals; it is frequently lethal for patients with chronic granulomatous disease (CGD), who carry mutations in the NADPH oxidase and thus cannot generate reactive oxygen species (ROS) in phagocytes^23,24^. Like other *Francisella* species, *F. philomiragia* is a facultative intracellular pathogen that escapes into the cytosol and replicates within host cells through the *Francisella* pathogenicity island-encoded type 6 secretion system^25^. However, unlike *Francisella novicida* or the highly virulent *Francisella tularensis*^26^, *F. philomiragia* is rapidly cleared by wild-type mice^18^. This striking resistance, combined with its intracellular lifestyle and the requirement of ROS for bacterial clearance^18^, suggested that *F. philomiragia* might provide a powerful model for identifying the innate immune mechanisms responsible for successful antibacterial immunity.

Here, we used a clinical isolate of *F. philomiragia* from a CGD patient^27–30^ (*Fp*^CGD^) to investigate how innate immune pathways coordinate bacterial clearance. We initially hypothesized that redundant cell death pathways cooperate with ROS to eliminate intracellular bacterial niches and clear the infection. Instead, we found that the protective function of inflammasome signaling is not cell death. Rather, NLRP3-derived IL-1β suppresses a detrimental RIPK3-dependent type I interferon response in mice deficient in caspase-8 and MLKL. This principle was validated in a second intracellular pathogen, *Listeria monocytogenes* lacking ActA-mediated cell-to-cell spread^31^, demonstrating that it is not unique to *Fp*^CGD^. In contrast, this necroptosis-independent role of RIPK3 protects against HSV-1 infection. These findings reveal an unexpected antagonism between inflammasome and RIPK3 signaling pathways and suggest that the outcome of infection is determined by the balance between competing antibacterial and antiviral immune programs.

## Results

### Mice deficient in inflammasomes, caspase-8, and MLKL are susceptible to *Fp*^CGD^

While *Francisella novicida* (*Fn*) is lethal to wild-type (WT) mice at doses as low as 10 CFU following intraperitoneal inoculation, WT mice were highly resistant to the closely related *Fp*^CGD^ (Figure 1A). WT mice rapidly cleared *Fp*^CGD^ within 24 hours and survived doses up to 10^7^ CFU (Figures S1A and 1A), consistent with it being an easily eliminated environmental pathogen. Consistent with the susceptibility of CGD patients^29^, mice deficient in ROS production (*Ncf1*^−/−^) were highly susceptible to *Fp*^CGD^ (Figures 1B and 1C)^18^, with 100 CFU proving lethal within 1 day post-infection (DPI) (Figure 1D). In contrast, reactive nitrogen species were dispensable for *Fp*^CGD^ control (Figures 1B and 1C). Further analysis demonstrated that adaptive immunity, complement (C3, C7), STING, and type I, II, and III interferons were all expendable for bacterial clearance (Figures S1B-S1J). Together, these findings suggest that *Fp*^CGD^ clearance relies on specific innate antibacterial mechanisms.

**Figure 1.**
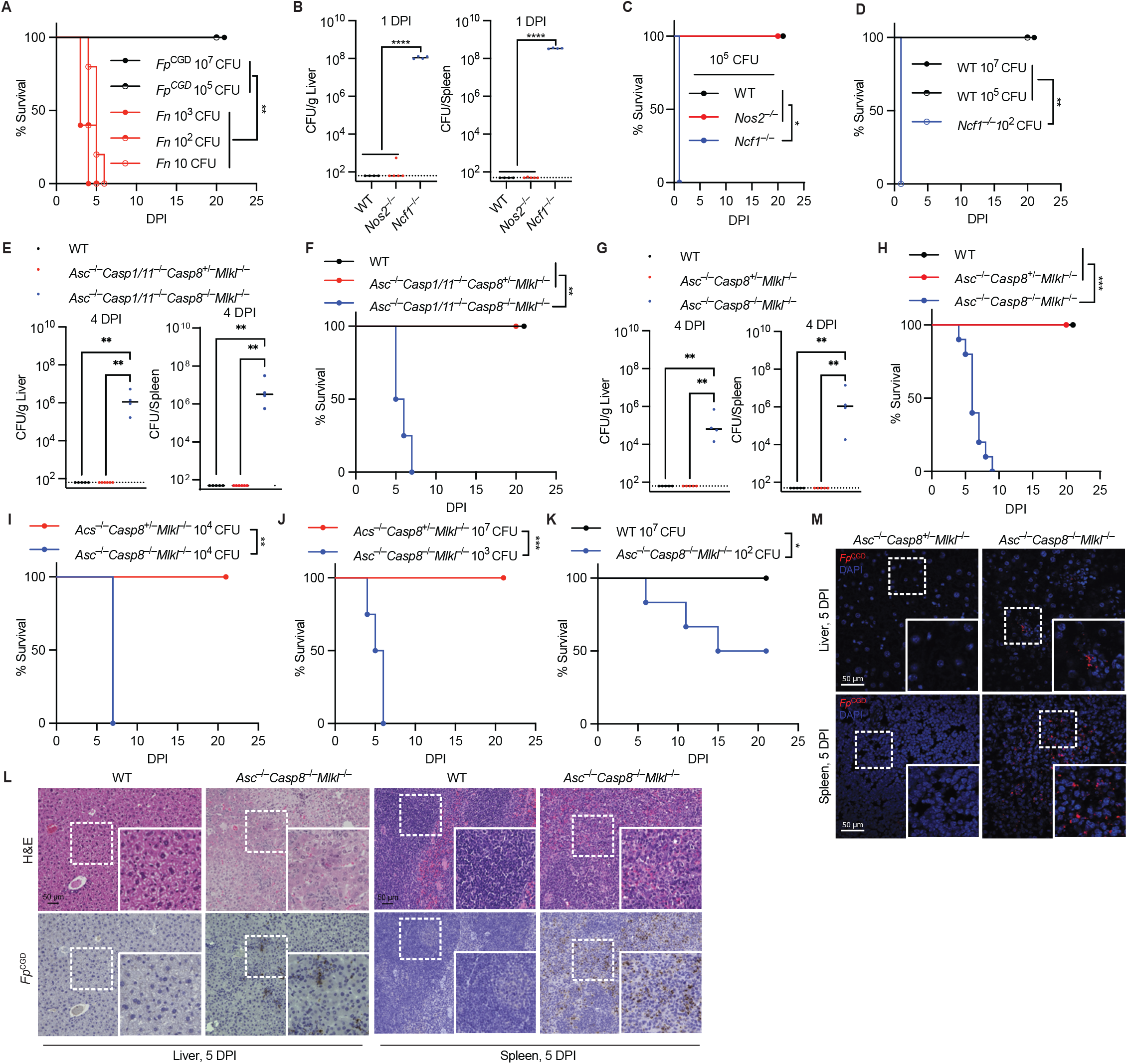
Combined loss of ASC, caspase-8, and MLKL causes susceptibility to *Fp*^CGD^. (A) WT mice were infected intraperitoneally (IP) with the indicated doses of *Fp*^CGD^ or *Fn,* and survival was monitored. Mice: n=5 for all. (B–M) Mice were infected IP with 10^5^ CFU of *Fp*^CGD^ unless otherwise indicated. CFU burdens in the liver and spleen were determined at the indicated time points (B, E, and G). Survival was monitored (C, D, F, and H–K). Livers and spleens were collected at 5 DPI for H&E staining and immunohistochemical (IHC) staining for *Fp*^CGD^ (brown) (L) or immunofluorescence staining for *Fp*^CGD^ (red) with DAPI (blue) (M). Mice: (B) WT=4, *Nos2*^−/−^=5, *Ncf1*^−/−^=4; (C) WT=4, *Nos2*^−/−^=3, *Ncf1*^−/−^=4; (D) n=5 for all; (E) WT=5, *Asc*^−/−^*Casp1/11*^−/−^*Casp8*^+/−^*Mlkl*^−/−^=6, *Asc*^−/−^*Casp1/11*^−/−^*Casp8*^−/−^*Mlkl*^−/−^=4; (F) WT=5, *Asc*^−/−^*Casp1/11*^−/−^*Casp8*^+/−^*Mlkl*^−/−^=5, *Asc*^−/−^*Casp1/11*^−/−^*Casp8*^−/−^*Mlkl*^−/−^=4; (G) WT=5, *Asc*^−/−^*Casp8*^+/−^*Mlkl*^−/−^=5, *Asc*^−/−^*Casp8*^−/−^*Mlkl*^−/−^=4; (H) WT=5, *Asc*^−/−^*Casp8*^+/−^*Mlkl*^−/−^=9, *Asc*^−/−^*Casp8*^−/−^*Mlkl*^−/−^=10; (I) *Asc*^−/−^*Casp8*^+/−^*Mlkl*^−/−^=6, *Asc*^−/−^*Casp8*^−/−^*Mlkl*^−/−^=3; (J) *Asc*^−/−^*Casp8*^+/−^*Mlkl*^−/−^=6, *Asc*^−/−^*Casp8*^−/−^*Mlkl*^−/−^=4; (K) WT=8, *Asc*^−/−^*Casp8*^−/−^*Mlkl*^−/−^=6; (L) n=4 for both; (M) n=3 for both. Data and statistical analyses are representative of at least two experiments (A, C, E, F, G, I, and J), or pooled from two experiments (B, D, H, and K). Dashed line indicates limit of detection. \**p* < 0.05; \*\**p* < 0.01; \*\*\**p* < 0.001; \*\*\*\**p* < 0.0001; survival curve analysis (A, C, D, F, and H–K), one-way ANOVA (B, E, and G). Scale bars: 50 μm in (L) and (M). See also Figure S1.

We previously showed that a flagellin-engineered *Salmonella* Typhimurium strain that triggers pyroptosis becomes trapped within pyroptotic macrophages, which are then efferocytosed by neutrophils and killed by ROS^32^. We therefore hypothesized that a similar pathway clears *Fp*^CGD^, where either pyroptosis, apoptosis, or necroptosis is required to promote efferocytosis by neutrophils and subsequent killing of bacteria by ROS. To test this hypothesis, we generated mice bearing compound deficiencies in multiple cell death pathways, beginning with *Asc*^−/−^*Casp1/11*^−/−^*Casp8*^−/−^*Mlkl*^−/−^ mice. In stark contrast to WT mice, these compound knockout mice displayed increased bacterial burdens in the liver and spleen and exhibited 100% mortality (Figures 1E and 1F). Restoring *Casp1/11* by generating *Asc*^−/−^*Casp8*^−/−^*Mlkl*^−/−^ mice did not alter the phenotype (Figures 1G-1H), confirming that caspase-11 is dispensable^17^ and that caspase-1 signaling arises through ASC. These phenotypes were potent, since susceptible mice fully succumbed to 10^4^ and 10^3^ CFU challenges and even showed 50% mortality after 10^2^ CFU challenges, whereas WT mice survived challenges of up to 10^7^ CFU (Figure 1I-1K). Infection occurred in small foci of immune cells scattered throughout both the liver and spleen, where bacteria could be easily identified by immunohistochemical or immunofluorescent staining (Figures 1L-1M). Collectively, these results demonstrate that mice deficient in inflammasome signaling, caspase-8, and MLKL are unable to control *Fp*^CGD^ infection.

### The NLRP3 inflammasome is activated by *Fp*^CGD^

We next sought to identify the inflammasome sensor that detects *Fp*^CGD^. We found that *Fp*^CGD^ specifically activates NLRP3 with negligible contribution from AIM2 in bone marrow-derived macrophages (BMMs), as evidenced by IL-1β processing and release, caspase-1 cleavage, propidium iodide (PI) uptake, and LDH release (Figures 2A-2D). *Fp*^CGD^-induced inflammasome activation required priming, such as by TLR stimulation with Pam3CSK4 or LPS (Figures 2A and S2A). ASC speck formation was readily observed in BMMs infected with *Fp*^CGD^, *Fn*, or a positive control (Figures 2E and 2F). We confirmed that, in contrast to *Fp*^CGD^, *Fn* activates only the AIM2 inflammasome^33,34^ (Figures 2D and S2B-S2D). As expected, these inflammasome signaling readouts were dependent on caspase-1 and GSDMD (Figures S2E-S2K). During *Fn* infection, prior publications indicate that cGAS first detects bacterial DNA, resulting in type I interferon production that primes guanylate-binding proteins that are required for AIM2 to detect *Fn*. We confirmed that *Fn*-induced inflammasome activation required cGAS; but in contrast, *Fp*^CGD^- induced inflammasome activation was cGAS-independent (Figures S2L-S2N)^34–36^.

**Figure 2.**
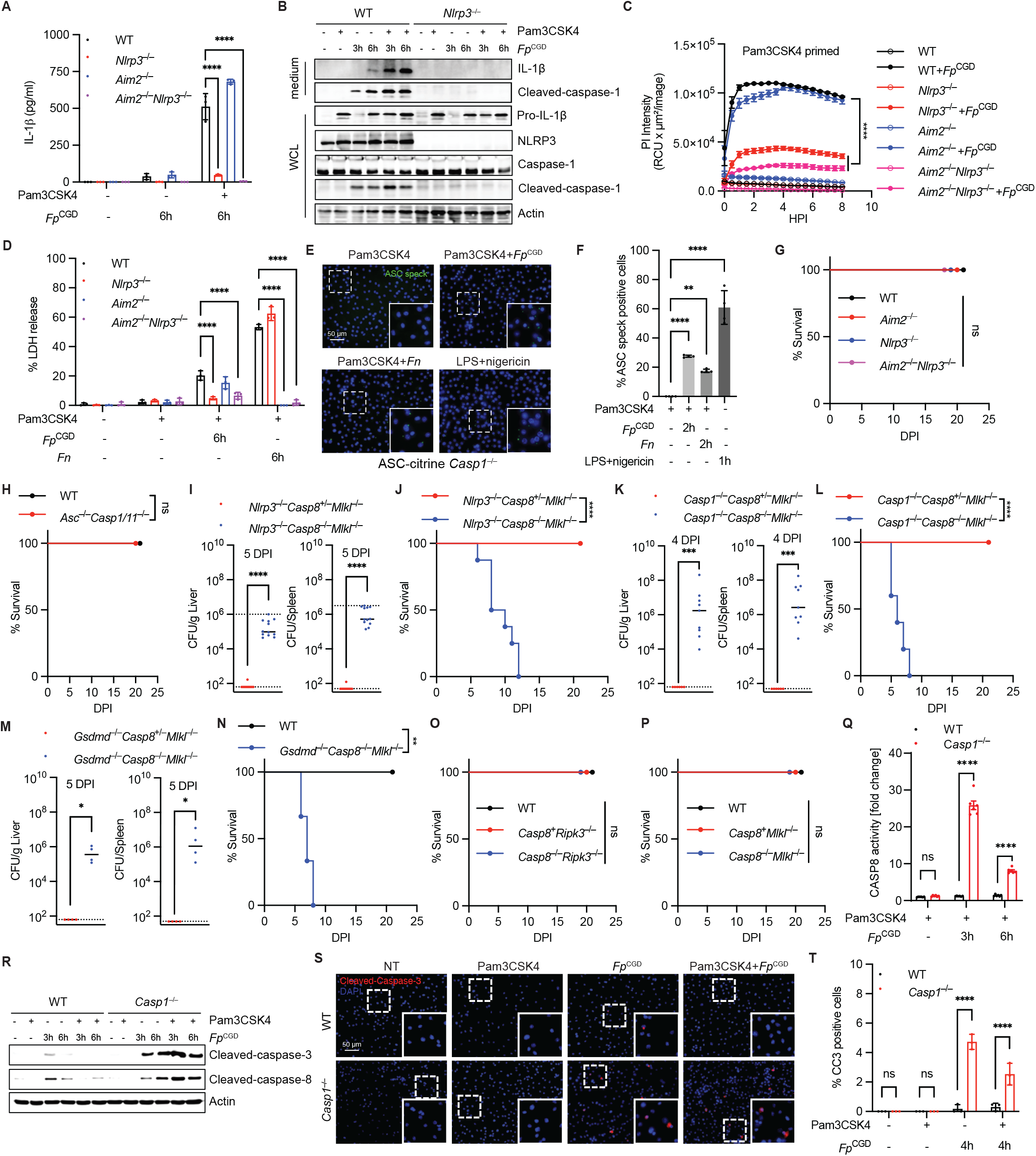
The NLRP3 pathway is required for resistance in compound knockout mice. (A– F) Bone marrow-derived macrophages (BMMs) were left unprimed or primed overnight with Pam3CSK4 (500 ng/mL) or LPS (100 ng/mL) and subsequently infected with *Fp*^CGD^ or *Fn* at an MOI of 5. IL-1β release was measured by ELISA at 6 hours post-infection (HPI) (A). Inflammasome activation was assessed by western blotting (B), PI uptake measured using IncuCyte (C), LDH release (D), and ASC speck formation at 2 HPI with quantification (E and F). LPS plus nigericin (10 μM) was used as a positive control for NLRP3 activation (E and F). (G–P) Mice were infected IP with 10^5^ CFU of *Fp*^CGD^. Survival was monitored (G, H, J, L, and N–P). CFU burdens in the liver and spleen were determined at the indicated time points (I, K, and M). Mice: (G and H) n=5 for all; (I) *Nlrp3*^−/−^*Casp8*^+/−^*Mlkl*^−/−^=10*, Nlrp3*^−/−^*Casp8*^−/−^*Mlkl*^−/−^=12; (J) *Nlrp3*^−/−^*Casp8*^+/−^*Mlkl*^−/−^=12*, Nlrp3*^−/−^*Casp8*^−/−^*Mlkl*^−/−^=8; (K) *Casp1*^−/−^*Casp8*^+/−^*Mlkl*^−/−^=6*, Casp1*^−/−^*Casp8*^−/−^*Mlkl*^−/−^=9; (L) *Casp1*^−/−^*Casp8*^+/−^*Mlkl*^−/−^=15*, Casp1*^−/−^*Casp8*^−/−^*Mlkl*^−/−^=5; (M) n=4 for both; (N) WT=5*, Gsdmd*^−/−^*Casp8*^−/−^*Mlkl*^−/−^=6; (O) WT=5*, Casp8*^+^*Ripk3*^−/−^=10, *Casp8*^−/−^*Ripk3*^−/−^=6; (P) WT=5*, Casp8*^+^*Ripk3*^−/−^=9, *Casp8*^−/−^*Ripk3*^−/−^=11. *Casp8*^+^ denotes both *Casp8*^+/+^ and *Casp8*^+/−^. (Q–T) BMMs were left unprimed or primed overnight with Pam3CSK4 (500 ng/mL) and infected with *Fp*^CGD^ at an MOI of 5. Apoptosis was assessed by caspase-8 activity assay (Q), western blotting (R), and immunofluorescence staining for cleaved caspase-3 (red) with quantification (S and T). Data and statistical analyses are representative of at least two experiments (A–F, H, and Q–T), pooled from two experiments (G, I–M, O, and P), or pooled from three experiments (N). Dashed line indicates limit of detection. ns, *p* > 0.05; \**p* < 0.05; \*\**p* < 0.01; \*\*\**p* < 0.001; \*\*\*\**p* < 0.0001; two-way ANOVA (A, D, Q, and T), one-way ANOVA (C and F), survival curve analysis (G, H, J, L, and N–P), and two-tailed unpaired t test (I, K, and M). Scale bars: 50 μm in (E) and (S). See also Figures S2–S4.

We further characterized *Fp*^CGD^-induced NLRP3 activation. Heat-killed *Fp*^CGD^ failed to induce IL- 1β or LDH release (Figures S2O and S2P). Potassium supplementation blocked *Fp*^CGD^-induced NLRP3 activation (Figure S2Q). ROS inhibition with N-acetylcysteine (NAC) had no effect on inflammasome activation (Figure S2R). These results are consistent with other known NLRP3 agonists^37,38^.

Despite robust NLRP3 inflammasome activation in vitro, mice deficient only in inflammasome genes mounted effective immune responses to *Fp*^CGD^ in vivo (Figures 2G, 2H, and S3A-S3L)^18^. Although *Asc*^−/−^ and *Nlrp3*^−/−^ mice showed slight increases in bacterial burden 16 hours post-infection (HPI) with high inocula (Figures S3K and S3L), these inflammasome-deficient mice ultimately survived the infection (Figures 2G and 2H), indicating that inflammasome signaling enhances control of *Fp*^CGD^, but that other innate defenses are sufficient in the absence of inflammasome defenses. Therefore, the importance of *Asc* only manifests in compound *Asc*^−/−^*Casp8*^−/−^*Mlkl*^−/−^ mice, which are highly susceptible.

We hypothesized that any gene in the inflammasome pathway could be substituted for the *Asc*^−/−^ allele. Consistent with this hypothesis, substituting the NLRP3 inhibitor MCC950 or a *Nlrp3*^−/−^ allele for *Asc*^−/−^ in the susceptible compound knockouts also resulted in susceptibility (Figures S3M, 2I and 2J). Downstream, we substituted the *Asc*^−/−^ allele with *Casp1*^−/−^ or *Gsdmd*^−/−^ alleles, and all the resulting triple knockout mice were equally susceptible (Figures 2K-2N). These findings indicate that the full NLRP3 pathway is required for protection in compound cell-death-deficient mice.

### Caspase-8 and necroptosis are dispensable for resistance to *Fp*^CGD^

Mice heterozygous for *Casp8* on the triple-knockout background were fully resistant to infection, indicating that susceptibility requires *Casp8* deletion (Figures 1E-1J, 1M, and 2I-2M). It remained possible that inflammasomes are dispensable and that *Casp8^−/−^Mlkl^−/−^* mice would be susceptible. To determine the individual contributions of caspase-8-mediated apoptosis and RIPK3-MLKL-mediated necroptosis, we examined single and double knockout mice. Single *Ripk3*^−/−^ or *Mlkl*^−/−^ mice were resistant to *Fp*^CGD^ (Figures S4A and S4B). Similarly, treatment with Nec-1s (a RIPK1 inhibitor) and infection of *Ripk3* kinase-dead mice had no adverse effect on bacterial clearance (Figures S4C-S4E). *Casp8*^−/−^*Ripk3*^−/−^ and *Casp8*^−/−^*Mlkl*^−/−^ mice had some detectable bacterial counts, but the phenotype was not strong enough to reach statistical significance (Figures S4F- S4I) and all these mice survived and cleared *Fp*^CGD^ (Figures 2O, 2P, S4J, and S4K). Furthermore, caspase-8 oligomerization-deficient (FGLG) and cleavage-resistant (D387A) mice, which cannot initiate apoptosis but retain caspase-8 scaffold and negative feedback functions^39^, were also resistant to *Fp*^CGD^ infection (Figures S4L and S4M). These results suggest that apoptosis and necroptosis are both dispensable for *Fp*^CGD^ clearance.

We next considered the hypothesis that pyroptotic, apoptotic, and necroptotic cell death act in triple redundancy; therefore, we examined BMMs in vitro for evidence of this. Inflammasome-deficient BMMs infected with *Fp*^CGD^ displayed enhanced caspase-8 and caspase-3 activation (Figures 2Q-2T, and S4N), indicating that BMMs failing to undergo pyroptosis are rerouted toward apoptosis, as expected^40,41^. These in vitro redundant cell death phenotypes might have explained the in vivo susceptibility of the compound cell death knockout mice; however, our next in vivo experiments would show this was not the case.

### RIPK3 becomes detrimental in the absence of the inflammasome, caspase-8, and MLKL

We found that inflammasome deletions are fully interchangeable in the susceptible triple knockout mice. Next, we examined whether the same was true for the necroptosis pathway. Strikingly, substitution of *Mlkl*^−/−^ with *Ripk3*^−/−^ in the triple knockouts resulted in opposite phenotypes. *Asc*^−/−^*Casp8*^−/−^*Ripk3*^−/−^ mice—unlike the susceptible *Asc*^−/−^*Casp8*^−/−^*Mlkl*^−/−^ strain—rapidly cleared *Fp*^CGD^ infection and all mice survived (Figures 3A-3C). To validate this unexpected finding, we substituted different inflammasome deficiencies onto the *Casp8*^−/−^*Ripk3*^−/−^ background. We generated *Casp1*^−/−^*Casp8*^−/−^*Ripk3*^−/−^ and *Asc*^−/−^*Casp1/11*^−/−^*Casp8*^−/−^*Ripk3*^−/−^ mice, both of which phenocopied the resistant phenotype of *Asc*^−/−^*Casp8*^−/−^*Ripk3*^−/−^ mice (Figures 3A, 3D-3G, and S5A). Notably, like WT mice, *Asc*^−/−^*Casp1/11*^−/−^*Casp8*^−/−^*Ripk3*^−/−^ mice survived a challenge of 10^7^ CFU (Figure 3H). Therefore, compound cell death knockout mice that include *Mlkl*^−/−^ succumb to *Fp*^CGD^ infection, whereas those that include *Ripk3*^−/−^ resist the infection.

**Figure 3.**
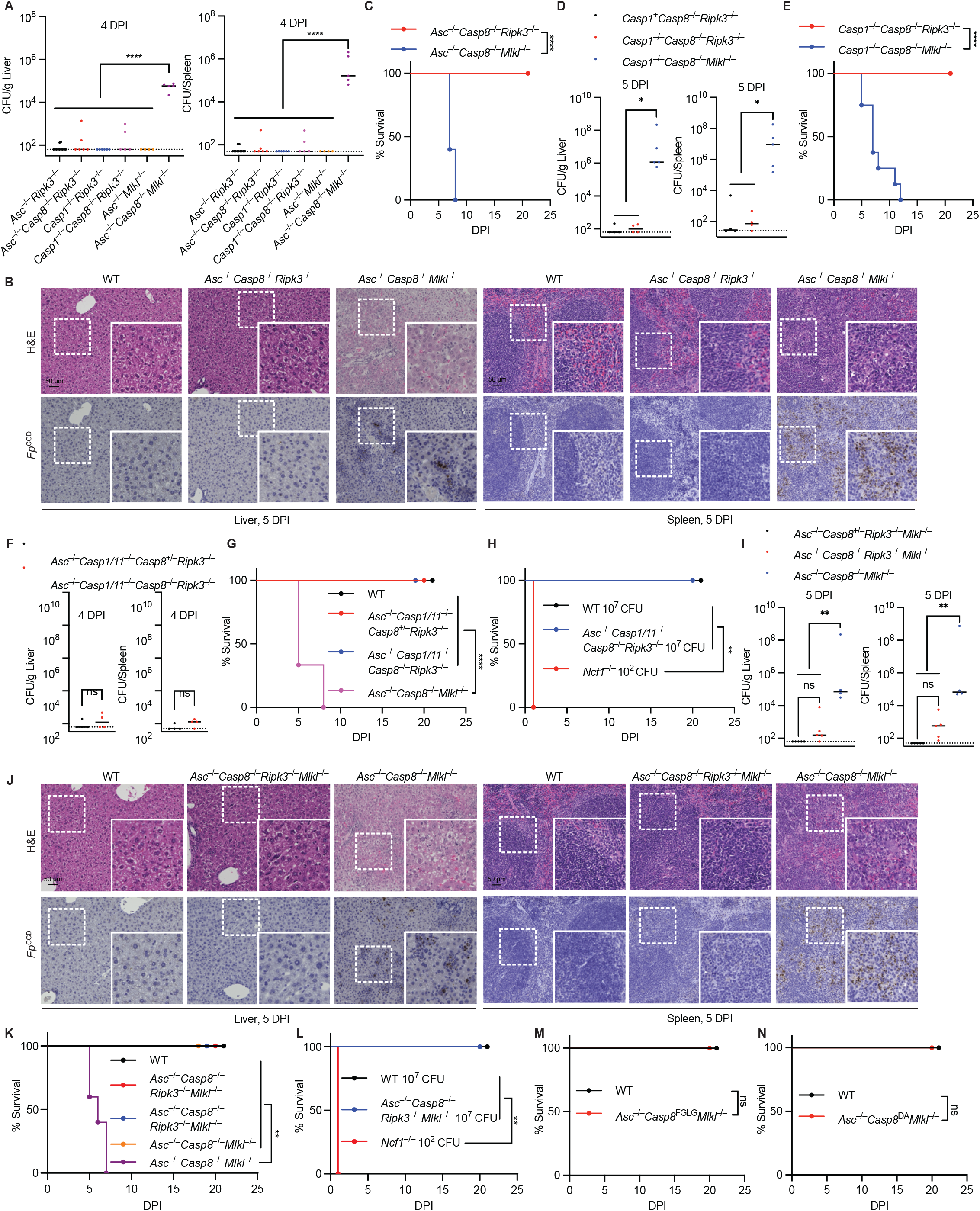
RIPK3 is detrimental during *Fp*^CGD^ infection in compound knockout mice. Mice were infected IP with 10^5^ CFU of *Fp*^CGD^ unless otherwise indicated. CFU burdens in the liver and spleen were determined at the indicated time points (A, D, F, and I). Survival was monitored (C, E, G, H, and K–N). Livers and spleens were collected at 5 DPI for H&E staining and IHC staining for *Fp*^CGD^ (B and J). Mice: (A) *Asc*^−/−^*Ripk3*^−/−^=11, *Asc*^−/−^*Casp8*^−/−^*Ripk3*^−/−^=6, *Casp1*^−/−^*Ripk3*^−/−^=6, *Casp1*^−/−^*Casp8*^−/−^*Ripk3*^−/−^=5, *Asc*^−/−^*Mlkl*^−/−^=4, *Asc*^−/−^*Casp8*^−/−^*Mlkl*^−/−^=5; (B) n=4 for all; (C) *Asc*^−/−^*Casp8*^−/−^*Ripk3*^−/−^=13, *Asc*^−/−^*Casp8*^−/−^*Mlkl*^−/−^=5; (D) *Casp1*^+^*Casp8*^−/−^*Ripk3*^−/−^=4, *Casp1*^−/−^*Casp8*^−/−^*Ripk3*^−/−^=4, *Casp1*^−/−^*Casp8*^−/−^*Mlkl*^−/−^=5, *Casp1*^+^ denotes both *Casp1*^+/+^ and *Casp1*^+/−^; (E) *Casp1*^−/−^*Casp8*^−/−^*Ripk3*^−/−^=9, *Casp1*^−/−^*Casp8*^−/−^*Mlkl*^−/−^=8; (F) n=4 for both; (G) WT=9, *Asc*^−/−^*Casp1/11*^−/−^*Casp8*^+/−^*Ripk3*^−/−^=6, *Asc*^−/−^*Casp1/11*^−/−^*Casp8*^−/−^*Ripk3*^−/−^=9, *Asc*^−/−^*Casp8*^−/−^*Mlkl*^−/−^=3; (H) n=5 for all; (I) *Asc*^−/−^*Casp8*^+/−^*Ripk3*^−/−^*Mlkl*^−/−^=5, *Asc*^−/−^*Casp8*^−/−^*Ripk3*^−/−^*Mlkl*^−/−^=5, *Asc*^−/−^*Casp8*^−/−^*Mlkl*^−/−^=4; (J) n=4 for all; (K) WT=5, *Asc*^−/−^*Casp8*^+/−^*Ripk3*^−/−^*Mlkl*^−/−^=5, *Asc*^−/−^*Casp8*^−/−^*Ripk3*^−/−^*Mlkl*^−/−^=6, *Asc*^−/−^*Casp8*^+/−^*Mlkl*^−/−^=7, *Asc*^−/−^*Casp8*^−/−^*Mlkl*^−/−^=5; (L) n=5 for all; (M) WT=8, *Asc*^−/−^*Casp8*^FGLG^*Mlkl*^−/−^=5; (N) WT=10, *Asc*^−/−^*Casp8*^DA^*Mlkl*^−/−^=9. Data and statistical analyses are pooled from two experiments (A, C, G, H, and K–N), representative of at least two experiments (D, F, and I), or pooled from three experiments (E). Dashed line indicates limit of detection. ns, *p* > 0.05; \**p* < 0.05; \*\**p* < 0.01; \*\*\*\**p* < 0.0001; one-way ANOVA (A, D, and I), survival curve analysis (C, E, G, H, and K–N), and two-tailed unpaired t test (F). Scale bars: 50 μm in (B) and (J). See also Figure S5.

These results could be explained by many possible mechanisms. We first considered the fact that *Casp8*^−/−^*Ripk3*^−/−^ mice develop lymphoproliferative disease (LPR) more slowly than *Casp8*^−/−^*Mlkl*^−/−^ mice^42^, but the severity catches up with time. For example, 20-week-old compound knockout mice carrying *Ripk3*^−/−^ had similar spleen weights to those of their 13-week-old *Mlkl*^−/−^ counterparts (Figure S5B), indicating roughly equivalent LPR. This normalization of the LPR by age had no effect on the phenotypes during *Fp*^CGD^ infection. Older compound knockout mice carrying *Ripk3*^−/−^ remained resistant (Figures S5C-S5E), and susceptible compound knockout mice carrying *Mlkl*^−/−^ could not be protected by infecting them at a young age before LPR becomes pronounced (Figures S5F-S5H). We conclude that LPR is not the primary driver of susceptibility.

Several publications have suggested that RIPK3 plays a role in driving transcriptional responses independently of MLKL^43–45^. If this pathway is active during *Fp*^CGD^ infection, it would mean that the susceptibility of *Asc*^−/−^*Casp8*^−/−^*Mlkl*^−/−^ mice is caused by the unrestrained presence of RIPK3. This hypothesis predicts that the genetic ablation of *Ripk3* in the susceptible triple knockout mice would protect them from *Fp*^CGD^ infection. Remarkably, *Asc*^−/−^*Casp8*^−/−^*Ripk3*^−/−^*Mlkl*^−/−^ mice became fully resistant to *Fp*^CGD^. These mice survived infection doses up to 10^7^ CFU and cleared bacteria from the spleen and liver (Figures 3I-3L, and S5I). These findings indicate that RIPK3 is actively detrimental in the susceptible compound cell death knockout mice.

Because the deletion of *Ripk3* protected the susceptible mice from infection, we can also conclude that pyroptosis, caspase-8-driven apoptosis, and necroptosis are all dispensable for defense against *Fp*^CGD^. Since cell death pathways per se are not required for defense, we hypothesized that the ability of caspase-8 to cleave and inactivate RIPK3 is critical^46^. To test this, we generated *Asc*^−/−^*Casp8*^FGLG^*Mlkl*^−/−^ and *Asc*^−/−^*Casp8*^DA^*Mlkl*^−/−^ mice, which express caspase-8 mutants that cannot initiate apoptosis but retain the ability to suppress RIPK3^39^. Both strains were fully resistant to *Fp*^CGD^ infection (Figures 3M, 3N, S5J, and S5K). Consistently, NLRP3 inhibition with MCC950 did not affect bacterial control in *Casp8*^FGLG^*Mlkl*^−/−^ mice (Figure S5L), further confirming that inflammasome-independent pathways drive this resistance. Therefore, in the absence of caspase-8-mediated cleavage, an unleashed RIPK3 performs a detrimental function.

### Cell death pathway requirements differ between pathogens

To assess the generalizability of our findings, we tested cell-death-deficient mice against commonly used host-adapted pathogens. *Salmonella enterica* serovar Typhimurium is an intracellular vacuolar bacterium that is highly lethal to C57BL/6 mice due to mutant *Nramp1* allele^47^. The Δ*aroA* mutant is used to reduce bacterial virulence, bringing the bacteria back into balance with C57BL/6 mice. The Δ*aroA* mutant causes a long-term infection that is eventually cleared within 7 to 12 weeks^48^. Consistent with a previous publication showing that redundant cell death pathways protect against the Δ*aroA* mutant^11^, *Asc*^−/−^*Casp8*^−/−^*Mlkl*^−/−^, *Asc*^−/−^*Casp8*^−/−^*Ripk3*^−/−^, *Asc*^−/−^*Casp8*^−/−^*Ripk3*^−/−^*Mlkl*^−/−^ and other compound cell-death-deficient mice were all uniformly susceptible to *Salmonella* Δ*aroA* (Figures 4A-4E). Similarly, these mice were susceptible to infection by *Listeria monocytogenes*, a cytosol-invasive bacterium (Figures 4F-4I, and S6A-S6F). Therefore, control of *S.* Typhimurium Δ*aroA* and *L. monocytogenes* appears to require functional cell death pathways.

**Figure 4.**
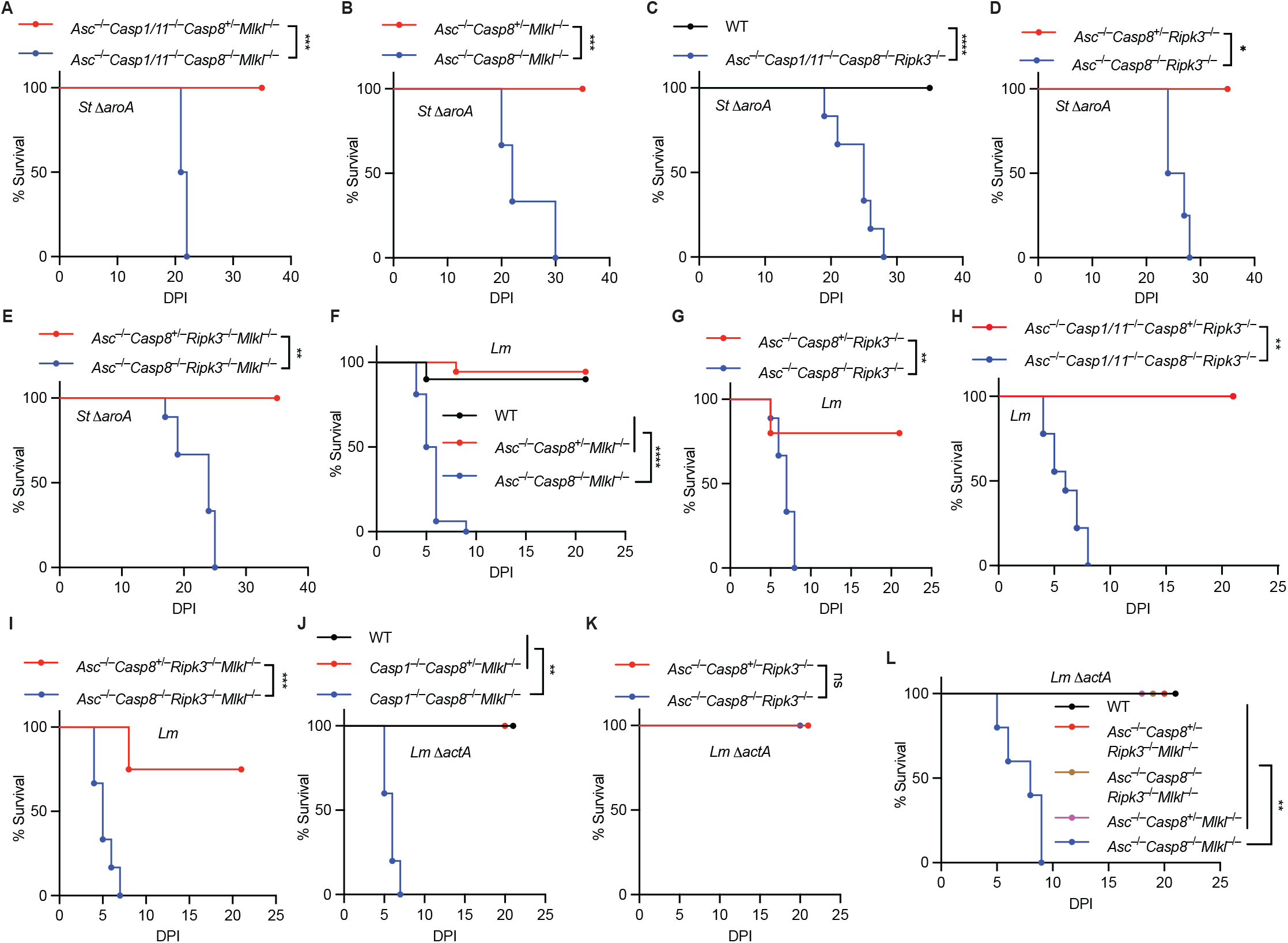
Host protection exhibits pathogen-specific requirements for cell death pathways. Mice were infected intravenously (IV) with *St* Δ*aroA* (2 × 10^2^ CFU; A–E), *Lm* (5 × 10^3^ CFU; F–I), or *Lm* Δ*actA* (1 × 10^7^ CFU; J–L). Survival was monitored. Mice: (A) *Asc*^−/−^*Casp1/11*^−/−^*Casp8*^+/−^*Mlkl*^−/−^=7, *Asc*^−/−^*Casp1/11*^−/−^*Casp8*^−/−^*Mlkl*^−/−^=6; (B) *Asc*^−/−^*Casp8*^+/−^*Mlkl*^−/−^=7, *Asc*^−/−^*Casp8*^−/−^*Mlkl*^−/−^=3; (C) WT=10, *Asc*^−/−^*Casp1/11*^−/−^*Casp8*^−/−^*Ripk3*^−/−^=6; (D) *Asc*^−/−^*Casp8*^+/−^*Ripk3*^−/−^=3, *Asc*^−/−^*Casp8*^−/−^*Ripk3*^−/−^=4; (E) *Asc*^−/−^*Casp8*^+/−^*Ripk3*^−/−^*Mlkl*^−/−^=4, *Asc*^−/−^*Casp8*^−/−^*Ripk3*^−/−^*Mlkl*^−/−^=9; (F) WT=10, *Asc*^−/−^*Casp8*^+/−^*Mlkl*^−/−^=18, *Asc*^−/−^*Casp8*^−/−^*Mlkl*^−/−^=16; (G) *Asc*^−/−^*Casp8*^+/−^*Ripk3*^−/−^=5, *Asc*^−/−^*Casp8*^−/−^*Ripk3*^−/−^=9; (H) *Asc*^−/−^*Casp1/11*^−/−^*Casp8*^+/−^*Ripk3*^−/−^=5, *Asc*^−/−^*Casp1/11*^−/−^*Casp8*^−/−^*Ripk3*^−/−^=9; (I) *Asc*^−/−^*Casp8*^+/−^*Ripk3*^−/−^*Mlkl*^−/−^=4, *Asc*^−/−^*Casp8*^−/−^*Ripk3*^−/−^*Mlkl*^−/−^=6; (J) n*=*5 for all; (K) n*=*4 for both; (L) WT=10, *Asc*^−/−^*Casp8*^+/−^*Ripk3*^−/−^*Mlkl*^−/−^=6, *Asc*^−/−^*Casp8*^−/−^*Ripk3*^−/−^*Mlkl*^−/−^=6; *Asc*^−/−^*Casp8*^+/−^*Mlkl*^−/−^=4, *Asc*^−/−^*Casp8*^−/−^*Mlkl*^−/−^=5. Data and statistical analyses are pooled from two experiments (A–E, G, and I–K), three experiments (F and L), or four experiments (H). ns, *p* > 0.05; \**p* < 0.05; \*\**p* < 0.01; \*\*\**p* < 0.001; \*\*\*\**p* < 0.0001; survival curve analysis for all. See also Figure S6.

Although *L. monocytogenes* and *Francisella* species are both cytosol-invasive, *L. monocytogenes* has the additional ability to spread from cell to cell using ActA to drive actin-based motility^49^. Therefore, Δ*actA L. monocytogenes* resembles *Francisella* species because both pathogens remain within the infected cell for longer periods of time. Strikingly, *L. monocytogenes* Δ*actA* adopted the same phenotypes as *Fp*^CGD^, in which compound knockout mice carrying *Mlkl*^−/−^ were highly susceptible, whereas those carrying *Ripk3*^−/−^ instead were completely resistant (Figures 4J and 4K). This phenotype was mirrored by bacterial burdens in the spleen, whereas liver burdens were comparable at 3 DPI (Figure S6G). Furthermore, as with *Fp*^CGD^, the addition of the *Ripk3*^−/−^ allele again protected the compound *Mlkl*^−/−^ mice (Figure 4L). This resistance pattern indicates that RIPK3-mediated susceptibility can be detrimental against both Gram-negative and Gram-positive bacterial pathogens that invade the cytosol with similar intracellular lifecycles.

### IL-1β antagonizes RIPK3-driven type I interferon signaling

Since cell death itself is dispensable for controlling *Fp*^CGD^, we reasoned that protective signals emanating from the inflammasome pathway antagonize the deleterious effects of RIPK3. IL-1β is an attractive candidate, as it mobilizes neutrophils^50^, the primary responders to bacterial infection and an abundant source of ROS. To test this hypothesis, we blocked IL-1β signaling in *Casp8*^−/−^*Mlkl*^−/−^ mice using anti-IL-1β antibodies, which resulted in significantly elevated bacterial burdens (Figure 5A). More definitively, *Il1b*^−/−^*Casp8*^−/−^*Mlkl*^−/−^ mice were highly susceptible to *Fp*^CGD^, exhibiting increased liver and spleen bacterial burdens and 100% mortality (Figures 5B and 5C). Conversely, recombinant IL-1β administration partially restored resistance in *Asc*^−/−^*Casp8*^−/−^*Mlkl*^−/−^ mice (Figure 5D). These results demonstrate that IL-1β is essential for counteracting RIPK3- mediated susceptibility.

**Figure 5.**
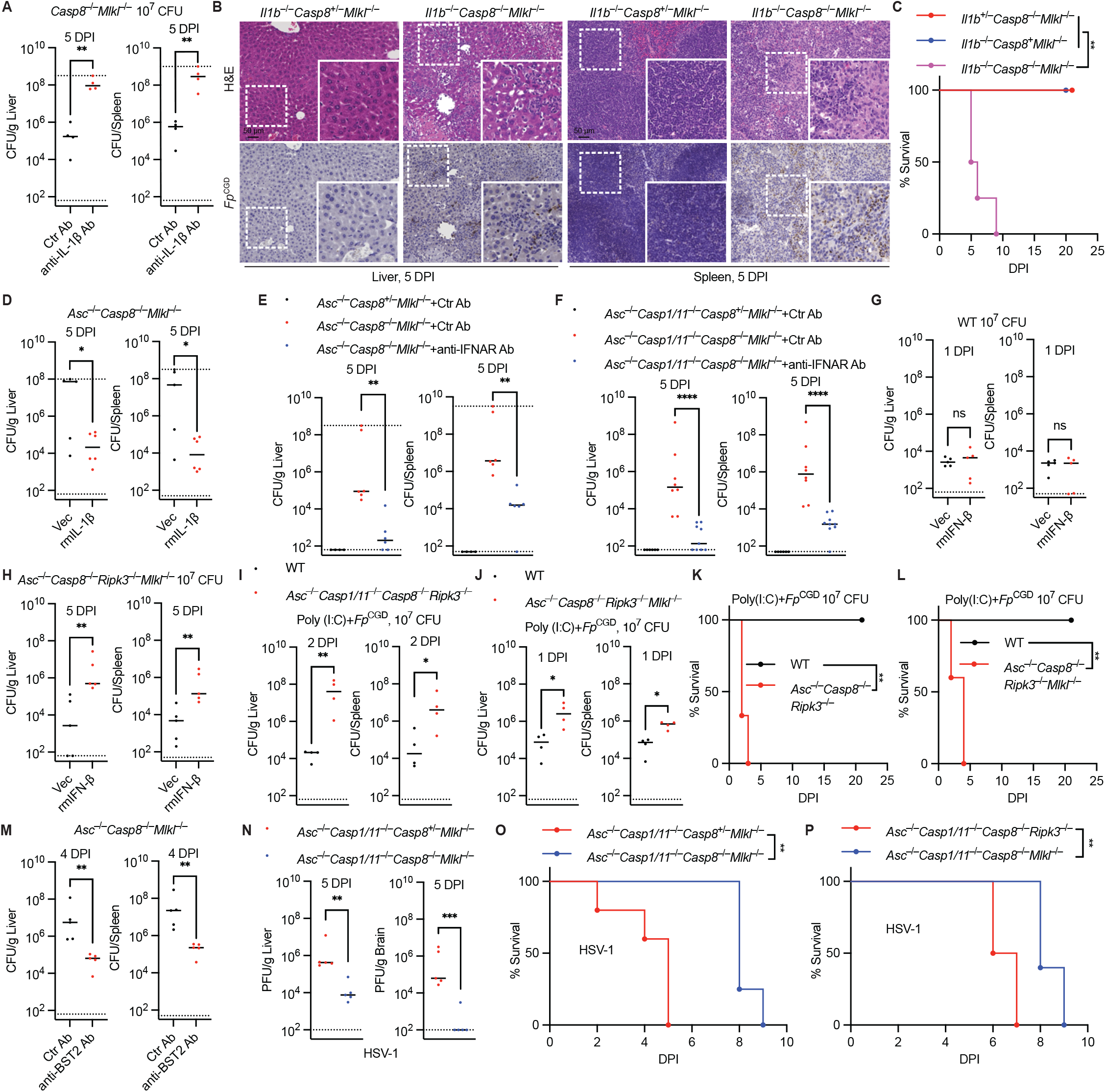
IL-1β counteracts RIPK3-driven type I interferon responses during *Fp*^CGD^ infection. (A–M) Mice were infected IP with 10^5^ CFU of *Fp*^CGD^ unless otherwise indicated. Mice received an isotype control antibody or anti-IL-1β antibody (0.2 mg/dose, IP, daily) beginning at −1 DPI, and CFU burdens were determined at 5 DPI (A). Livers and spleens were collected at 5 DPI for H&E staining and IHC staining for *Fp*^CGD^ (B). Survival was monitored (C). Mice were treated with vehicle or recombinant mouse IL-1β (1 μg/mouse, IP, daily) beginning at −3 DPI, and CFU burdens were determined at 5 DPI (D). Mice received an isotype control antibody or anti-IFNAR antibody (0.2 mg/dose, IP, daily) beginning at −1 DPI, and CFU burdens were determined at 5 DPI (E and F). Mice were treated with vehicle or recombinant mouse IFN-β (100 ng/mouse, IP, daily) beginning at −1 DPI, and CFU burdens were determined at the indicated time points (G and H). Mice were treated with low-molecular-weight poly(I:C) (5 mg/kg, IP, daily) from −1 to 5 DPI and CFU burdens were determined at the indicated time points (I and J) or survival was monitored (K and L). Mice received an isotype control antibody or anti-BST2 antibody (0.2 mg/dose, IP, daily) beginning at −1 DPI, and CFU burdens were determined at 4 DPI (M). Mice: (A–C) n=4 for all, *Casp8*^+^ denotes both *Casp8*^+/+^ and *Casp8*^+/−^; (D) Vec=5, rmIL-1β=6; (E) *Asc*^−/−^*Casp8*^+/−^*Mlkl*^−/−^+Ctr Ab=4, *Asc*^−/−^*Casp8*^−/−^*Mlkl*^−/−^+Ctr Ab=6, *Asc*^−/−^*Casp8*^−/−^*Mlkl*^−/−^+anti-IFNAR Ab=6; (F) *Asc*^−/−^*Casp1/11*^−/−^*Casp8*^+/−^*Mlkl*^−/−^+Ctr Ab=6, *Asc*^−/−^*Casp1/11*^−/−^*Casp8*^−/−^*Mlkl*^−/−^+Ctr Ab=8, *Asc*^−/−^*Casp1/11*^−/−^*Casp8*^−/−^*Mlkl*^−/−^+anti-IFNAR Ab=9; (G and H) n=5 for all; (I and J) n=4 for all; (K) WT=5, *Asc*^−/−^*Casp8*^−/−^*Ripk3*^−/−^=6; (L and M) n=5 for all. (N–P) Mice were infected IP with HSV-1 strain SC16 (5 × 10^6^ PFU) and viral titers in the liver and brain were determined at 5 DPI (N) or survival was monitored (O and P). Mice: (N) n=5 for both; (O) *Asc*^−/−^*Casp1/11*^−/−^*Casp8*^+/−^*Mlkl*^−/−^=5, *Asc*^−/−^*Casp1/11*^−/−^*Casp8*^−/−^*Mlkl*^−/−^=4; (P) *Asc*^−/−^*Casp1/11*^−/−^*Casp8*^−/−^*Ripk3*^−/−^=8, *Asc*^−/−^*Casp1/11*^−/−^*Casp8*^−/−^*Mlkl*^−/−^=5. Data and statistical analyses are representative of at least two experiments (A, I, and J), pooled from two experiments (C–E, G, H, K–P), or pooled from three experiments (F). Dashed line indicates limit of detection. ns, *p* > 0.05; \**p* < 0.05; \*\**p* < 0.01; \*\*\**p* < 0.001; \*\*\*\**p* < 0.0001; two-tailed unpaired t test (A, D, G–J, M, and N), survival curve analysis (C, K, L, O, and P), and one-way ANOVA (E and F). Scale bars: 50 μm in (B). See also Figure S7.

We next investigated the mechanism underlying the detrimental function of RIPK3. The literature suggests that RIPK3 drives the production of proinflammatory cytokines, including type I interferons^43–45^, which have been shown to exacerbate certain bacterial infections^51^. To test whether RIPK3-driven type I interferon production is responsible for *Fp*^CGD^ susceptibility, we blocked IFNAR signaling in susceptible compound knockout mice. Remarkably, IFNAR blockade substantially protected these mice from *Fp*^CGD^ infection (Figures 5E and 5F). Conversely, exogenous type I interferon administration caused the otherwise-resistant compound knockout mice to become susceptible to infection (Figure 5H), whereas type I interferon treatment had minimal effect on WT mice (Figure 5G). Consistent with these findings, poly(I:C) treatment (a TLR3 agonist that drives type I interferon production) likewise sensitized compound knockout mice carrying *Ripk3*^−/−^ (Figures 5I-5L), further supporting the hypothesis that RIPK3-driven type I interferon is detrimental. Resistant mice do not require interferon responses to clear *Fp*^CGD^, as blocking both IFNAR and IFNGR pathways did not impair *Fp*^CGD^ clearance in resistant compound knockout mice (Figure S7A). Furthermore, as previously noted, single and compound interferon receptor knockout mice remain resistant (Figures S1F-S1J). A recent publication suggests that the RHIM domain of RIPK3 drives its transcriptional response^45^. Consistent with this, blocking RIPK3 kinase activity in susceptible compound knockout mice had no effect on their susceptibility to *Fp*^CGD^ (Figure S7B). This suggests that the scaffolding function of RIPK3, rather than its kinase activity, mediates this susceptibility.

We considered which cell type might be the source of this detrimental type I interferon. Plasmacytoid dendritic cells (pDCs) are a low-abundance cell type that produces copious amounts of type I interferon during viral infection^52^. We therefore depleted pDCs in susceptible compound knockout mice using anti-BST2 antibodies. Intriguingly, pDC depletion substantially protected the susceptible mice (Figures 5M and S7C), suggesting that pDC-derived type I interferon contributes to RIPK3-driven immunopathology. This protection was not complete, which could be due to incomplete depletion or contributions from other sources of type I interferon. These results indicate that pDCs, and perhaps other cell types, experience a maladaptive RIPK3 signaling event during *Fp*^CGD^ infection. This aberrant type I interferon response is driven by genetic caspase-8 deficiency, rather than by direct inhibition of caspase-8 activity by a *Fp*^CGD^ virulence factor.

### RIPK3-driven responses are beneficial against viral infection

In contrast to *Fp*^CGD^, herpesviruses actively inhibit caspase-8 in infected cells^53^. Since type I interferons provide protective immunity against viral infections^54^, we hypothesized that the RIPK3- driven type I interferon response would be protective during viral challenge. We infected *Asc*^−/−^*Casp1/11*^−/−^*Casp8*^−/−^*Mlkl*^−/−^ mice (which are sufficient for RIPK3) with HSV-1 and compared outcomes to littermate controls. Intriguingly, these compound knockout mice exhibited reduced viral titers and prolonged survival compared to their control littermates (Figures 5N, 5O, and S7D). Most importantly, compound knockout mice carrying *Mlkl*^−/−^ survived longer than those carrying *Ripk3*^−/−^ (Figure 5P), suggesting that RIPK3-mediated responses are protective against herpes simplex virus via an MLKL-independent pathway. The contrasting outcomes during HSV-1 and *Fp*^CGD^ infection indicate that the immune consequences of RIPK3 activation are detrimental during bacterial infection but protective during viral infection.

### Neutrophils are essential for resistance to *Fp*^CGD^ whereas macrophages provide a replicative niche

Macrophages and neutrophils are the primary cell types producing ROS via the NADPH oxidase, with neutrophils producing far more ROS than macrophages^55,56^. To validate that ROS production arises from hematopoietic cells, we reconstituted lethally irradiated WT mice with bone marrow from *Ncf1*^−/−^ donors (lacking ROS production). The resulting chimeric mice were highly susceptible to *Fp*^CGD^ infection. In contrast, *Ncf1*^−/−^ mice receiving WT bone marrow were completely protected (Figure 6A).

**Figure 6.**
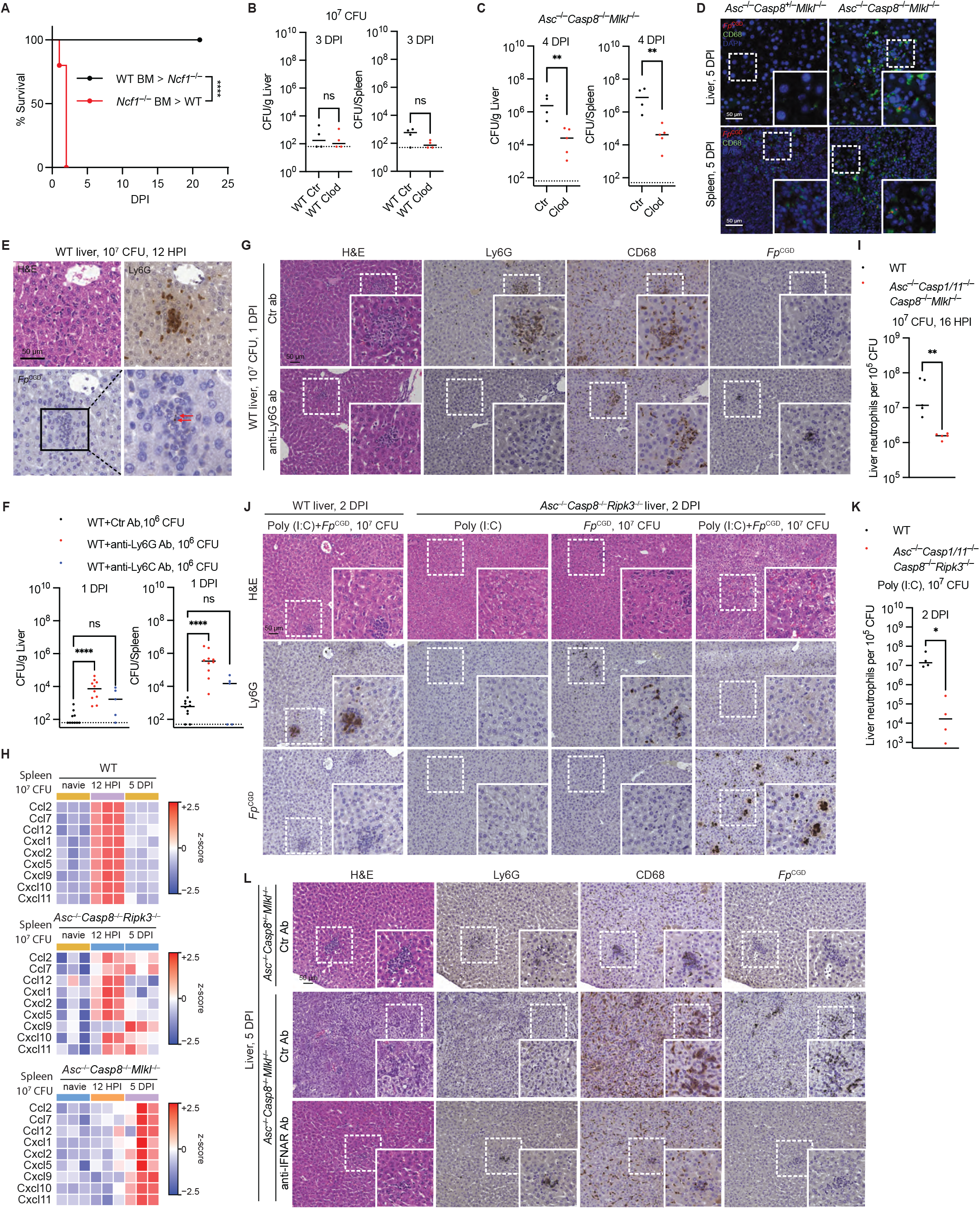
Type I interferon suppresses neutrophil responses during *Fp*^CGD^ infection. (A–G) Mice were infected IP with 10^5^ CFU of *Fp*^CGD^ unless otherwise indicated. Survival was monitored (A). Mice received control liposomes or clodronate liposomes (200 μL, IV, every other day) beginning at −1 DPI, and CFU burdens were determined at the indicated time points (B and C). Livers and spleens were collected at 5 DPI for immunofluorescence staining for *Fp*^CGD^ (red) and CD68 (green) (D). Livers were collected at 12 HPI for H&E staining and IHC staining for Ly6G and *Fp*^CGD^ (E). Mice received isotype control antibody, anti-Ly6G antibody, or anti-Ly6C antibody (0.2 mg/dose, IP, daily) beginning at −1 DPI, and CFU burdens were determined at 1 DPI (F). Mice received isotype control antibody or anti-Ly6G antibody (0.2 mg/dose, IP, daily) beginning at −1 DPI. Livers were collected at 1 DPI for H&E staining and IHC staining for Ly6G, CD68, and *Fp*^CGD^ (G). (H–K) Mice were infected with 10^7^ CFU of *Fp*^CGD^. Spleens from naïve mice, as well as infected mice at 12 HPI and 5 DPI, were snap-frozen for RNA extraction using TRIzol, followed by bulk RNA sequencing. Chemokine expression among the top 50 induced genes is shown (H). Livers were collected at the indicated time points for flow cytometry to determine the efficiency of neutrophil recruitment (I and K). Mice were treated with low-molecular-weight poly(I:C) (5 mg/kg, IP, daily) from −1 to 1 DPI. Livers were harvested at 2 DPI for H&E staining and IHC staining for Ly6G and *Fp*^CGD^ (J). (L) Mice received either an isotype control antibody or an anti-IFNAR antibody (0.2 mg/dose, IP, daily) beginning at −1 DPI. Mice were infected IP with 10^5^ CFU of *Fp*^CGD^. Livers were collected at 5 DPI for H&E staining and IHC staining of Ly6G, CD68, and *Fp*^CGD^. Mice: (A) WT BM>*Ncf1*^−/−^=11, *Ncf1*^−/−^ BM>WT=10; (B) n=4 for both; (C) Ctr=4, Clod=5; (D and E) n=3 for all; (F) Ctr Ab=10, anti-Ly6G Ab=10, anti-Ly6C Ab=5; (G) n=3 for both; (H) n=3–6 for each group; (I) n=5 for both; (J–L) n=4 for all. Data and statistical analyses are pooled from two experiments (A, C, and F) or representative of at least two experiments (B). Dashed line indicates limit of detection. ns, *p* > 0.05; \**p* < 0.05; \*\**p* < 0.01; \*\*\**p* < 0.001; \*\*\*\**p* < 0.0001; survival curve analysis (A), two-tailed unpaired t test (B, C, I, and K), and one-way ANOVA (F). Scale bars: 50 μm in (D), (E), (G), (J), and (L). See also Figure S8.

To test the role of macrophages, we depleted them in WT mice using clodronate liposomes, which surprisingly had no effect on bacterial clearance (Figure 6B). *Ccr2*^−/−^ mice, which are deficient in monocyte recruitment^22,57^, were also resistant to *Fp*^CGD^ infection (Figures S8A and S8B), further supporting the dispensable role of macrophages in WT mice. Interestingly, macrophage depletion substantially protected susceptible compound knockout mice (Figures 6C and S8C). Moreover, CD68⁺ macrophages colocalized with *Fp*^CGD^ in the liver and spleen of these susceptible mice (Figure 6D), suggesting that macrophages may serve as replicative niches for the bacteria, which is similar to other *Francisella* species.

We hypothesized that neutrophils are the primary cells that kill *Fp*^CGD^. Immunohistochemistry and immunofluorescence microscopy revealed that neutrophils were recruited to the livers of WT mice and colocalized with *Fp*^CGD^ (Figures 6E and S8D), suggesting their role as primary responders to infection. Next, we depleted neutrophils in WT mice using an anti-Ly6G antibody (clone 1A8). At 1 DPI, neutrophil-depleted mice exhibited significantly elevated bacterial burdens in the liver and spleen compared to controls (Figures 6F, 6G, and S8E). In contrast, anti-Ly6C antibody-mediated depletion of inflammatory monocytes had no significant effect (Figure 6F). In control WT mice, both Ly6G⁺ neutrophils and CD68⁺ macrophages were recruited to liver infection loci, leading to minimal *Fp*^CGD^ staining (Figure 6G). In anti-Ly6G-treated WT mice, CD68⁺ macrophages predominated at these loci, yet significant *Fp*^CGD^ staining remained (Figure 6G). Of note, resident liver macrophages—Kupffer cells—are also CD68⁺. This rapid increase in bacterial burden within 24 hours of neutrophil depletion demonstrates that neutrophils are essential for bacterial control. Altogether, these findings demonstrate that ROS production within the neutrophil compartment clears the infection.

### Type I interferon signaling suppresses neutrophil response

We next sought to define how excessive type I interferon signaling disrupts antibacterial immunity. We hypothesized that WT and resistant *Asc*^−/−^*Casp8*^−/−^*Ripk3*^−/−^ mice mount an appropriate early transcriptional response to infection, whereas susceptible *Asc*^−/−^*Casp8*^−/−^*Mlkl*^−/−^ mice fail to initiate this protective program.

To test this hypothesis in an unbiased manner, we performed bulk RNA sequencing on spleens collected from naïve mice, at an early stage of infection (12 HPI), before bacterial burdens diverged, and at 5 DPI, when resistant mice had largely cleared the infection whereas susceptible mice carried significant bacterial burdens. In WT spleens, the 50 most highly induced genes at 12 HPI were dominated by two functional programs (Figures S8F and 6H). First, multiple chemokines involved in leukocyte recruitment were strongly induced, including CXCR1/2 ligands that recruit neutrophils (*Cxcl1*, *Cxcl2*, and *Cxcl5*), CCR2 ligands that recruit inflammatory monocytes (*Ccl2*, *Ccl7*, and *Ccl12*), and CXCR3 ligands that recruit type 1 lymphocytes (*Cxcl9*, *Cxcl10*, and *Cxcl11*). Second, numerous interferon-stimulated genes (ISGs), including *Gbp2*, *Gbp4*, *Gbp5*, *Socs1*, and *Slfn4*, were induced, together with the ISG chemokines *Cxcl9*, *Cxcl10*, and *Cxcl11*. By 5 DPI, this inflammatory signature had largely resolved in wild-type mice, consistent with successful bacterial clearance.

Resistant *Asc*^−/−^*Casp8*^−/−^*Ripk3*^−/−^ mice mounted a remarkably similar transcriptional response, although the magnitude of induction was modestly reduced and inflammatory gene expression had not completely returned to baseline by 5 DPI (Figures 6H), consistent with their delayed bacterial clearance. In striking contrast, susceptible *Asc*^−/−^*Casp8*^−/−^*Mlkl*^−/−^ mice failed to induce the chemokine program at 12 HPI. By 5 DPI, when bacterial burdens had become massive, these mice finally exhibited robust chemokine expression (Figures 6H). Thus, susceptible mice do not fail to activate inflammatory pathways altogether; rather, they mount the correct response too late to prevent uncontrolled bacterial replication.

Because RIPK3-driven type I interferon signaling promoted susceptibility, and type I interferons have been reported to suppress neutrophil responses during other bacterial infections^51^, we next investigated whether neutrophil recruitment was impaired in the susceptible compound knockout mice. Consistent with the transcriptional data, susceptible compound knockout mice were less efficient at recruiting neutrophils to the liver during the early phase of infection than resistant mice (Figures S8G and 6I). This observation suggested that excessive type I interferon signaling suppresses the early neutrophil response required for bacterial control.

To determine whether elevated type I interferon signaling is sufficient to impair neutrophil recruitment, we treated otherwise resistant *Asc*^−/−^*Casp8*^−/−^*Ripk3*^−/−^ and *Asc*^−/−^*Casp1/11*^−/−^*Casp8*^−/−^*Ripk3*^−/−^ mice with poly(I:C). Poly(I:C) markedly increased bacterial burdens while largely abolishing liver neutrophil recruitment (Figures 6J and 6K). In contrast, WT mice remained resistant despite poly(I:C) treatment, suggesting that inflammasome activation counteracts the detrimental effects of excessive type I interferon signaling (Figure 6J). Importantly, poly(I:C) alone did not cause gross tissue pathology in *Asc*^−/−^*Casp8*^−/−^*Ripk3*^−/−^ mice (Figure 6J), indicating that susceptibility specifically results from the combination of bacterial infection and excessive type I interferon signaling.

Next, we asked whether restoring resistance by blocking type I interferon signaling also restored neutrophil responses in susceptible compound knockout mice. Histological analysis at 5 DPI showed that livers from isotype control-treated *Asc*^−/−^*Casp8*^−/−^*Mlkl*^−/−^ mice contained extensive bacterial aggregates that colocalized predominantly with macrophages, with relatively few neutrophils (Figure 6L), consistent with macrophages serving as bacterial replication niches. Notably, livers from IFNAR-blocked *Asc*^−/−^*Casp8*^−/−^*Mlkl*^−/−^ mice had largely returned to tissue homeostasis by 5 DPI (Figure S8H), consistent with the resolution of the infection. The few remaining inflammatory foci contained abundant neutrophils and macrophages but little detectable *Fp*^CGD^ staining, resembling resistant *Asc*^−/−^*Casp8*^+/−^*Mlkl*^−/−^ mice (Figure 6L). Therefore, consistent with the marked reduction in bacterial burdens, IFNAR blockade also restored neutrophil accumulation within infected tissues.

Likewise, spleens from isotype control-treated *Asc*^−/−^*Casp8*^−/−^*Mlkl*^−/−^ mice displayed extensive *Fp*^CGD^ staining, whereas IFNAR-blocked *Asc*^−/−^*Casp8*^−/−^*Mlkl*^−/−^ mice showed significant clearance of the infection (Figure S8I). Together, these findings demonstrate that excessive type I interferon signaling suppresses the timely recruitment of neutrophils required for effective antibacterial immunity, thereby permitting uncontrolled bacterial replication and disease progression.

### NLRP3 activation in neutrophils is caspase-8-independent

An apparent paradox emerged: *Casp8*^−/−^*Mlkl*^−/−^ mice were resistant to *Fp*^CGD^ (Figures 2P), whereas *Nlrp3*^−/−^*Casp8*^−/−^*Mlkl*^−/−^ mice were susceptible (Figures 2I and 2J), indicating that NLRP3 inflammasome signaling remains functional in the absence of caspase-8. However, published studies demonstrate that caspase-8 is required for NLRP3 inflammasome activation in macrophages in vitro^58,59^. We verified that this phenotype held true during *Fp*^CGD^ infection, as neither *Casp8*^−/−^*Mlkl*^−/−^ nor *Casp8*^−/−^*Ripk3*^−/−^ BMMs activated NLRP3 in response to *Fp*^CGD^ infection, even after priming with TLR2 or TLR4 ligands (Figures 7A-7C and S9A-S9K). These double knockouts also expressed lower levels of pro-IL-1β (Figures 7B, S9E, S9J, and S9K). In contrast, *Fn*-induced inflammasome activation in BMMs was independent of caspase-8 (Figures S9L-S9N), suggesting that AIM2 activation is unaffected in caspase-8-deficient BMMs. Altogether, these results suggest that pro-IL-1β expression and NLRP3 activation could occur in vivo in cell types other than macrophages in caspase-8-deficient mice.

**Figure 7.**
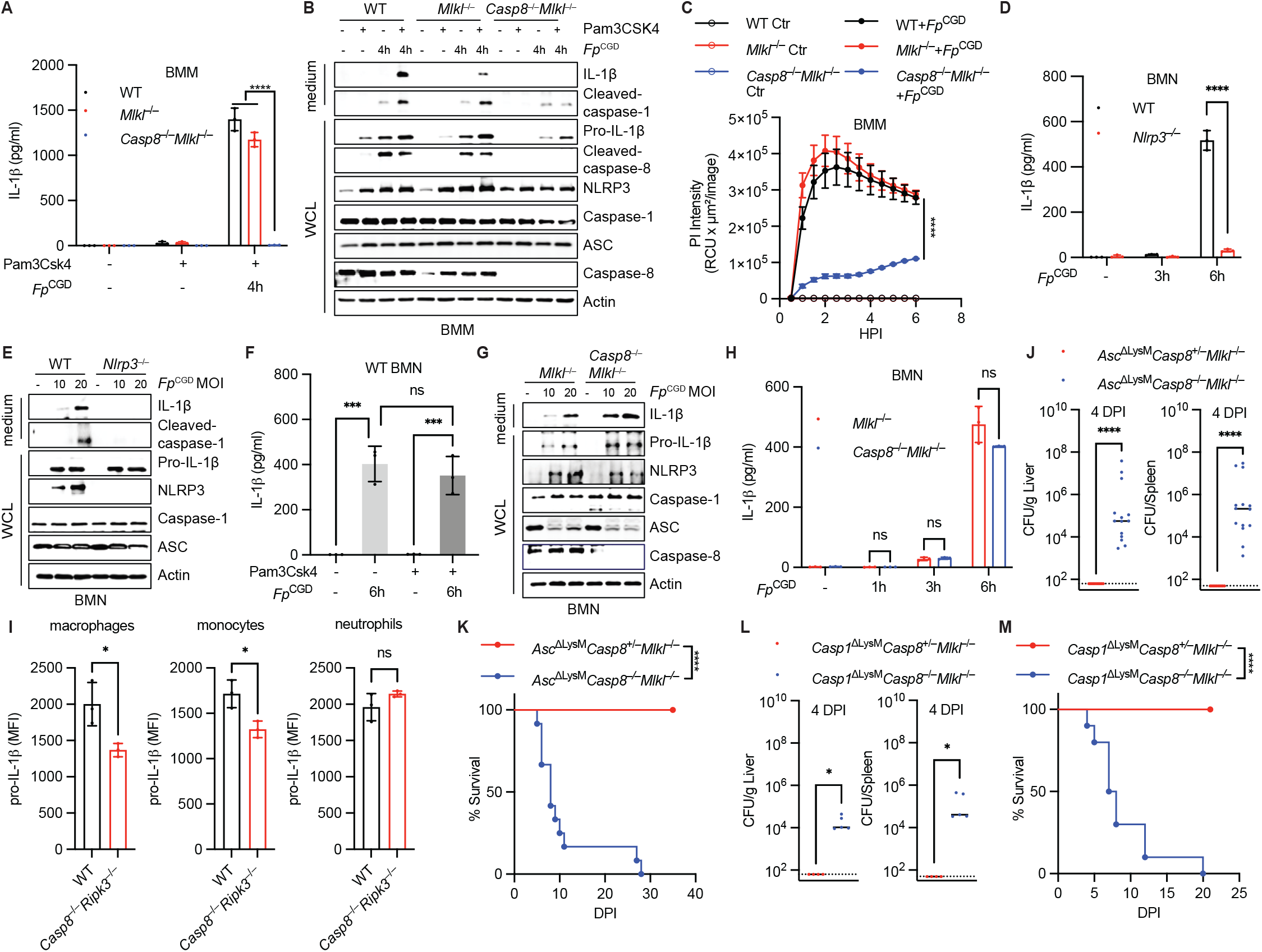
Inflammasome signaling in myeloid cells is protective against *Fp*^CGD^. (A–C) BMMs were left unprimed or primed overnight with Pam3CSK4 (500 ng/mL) and infected with *Fp*^CGD^ at an MOI of 5. IL-1β release was measured by ELISA (A), inflammasome activation was assessed by western blotting (B), and PI uptake was measured using IncuCyte (C). (D–H) Bone marrow-derived neutrophils (BMNs) were infected with *Fp*^CGD^ at an MOI of 20 (D–G) or 10 (H). Culture supernatants were collected at the indicated time points (D and H) or at 6 HPI (F) for IL-1β measurement by ELISA. Western blotting was performed at 6 HPI (E and G). (I) Mice were infected IP with 10^7^ CFU of *Fp*^CGD^, and livers were harvested at 16 HPI for flow cytometric analysis. (J–M) Mice were infected IP with 10^5^ CFU of *Fp*^CGD^. CFU burdens in the liver and spleen were determined at 4 DPI (J and L). Survival was monitored (K and M). Mice: (I) n=3 for both; (J) *Asc*^ΔLysM^*Casp8*^+/−^*Mlkl*^−/−^=10, *Asc*^ΔLysM^*Casp8*^−/−^*Mlkl*^−/−^=14; (K) *Asc*^ΔLysM^*Casp8*^+/−^*Mlkl*^−/−^=14, *Asc*^ΔLysM^*Casp8*^−/−^*Mlkl*^−/−^=12; (L) *Casp1*^ΔLysM^*Casp8*^+/−^*Mlkl*^−/−^=4, *Casp1*^ΔLysM^*Casp8*^−/−^*Mlkl*^−/−^=5; (M) *Casp1*^ΔLysM^*Casp8*^+/−^*Mlkl*^−/−^=9, *Casp1*^ΔLysM^*Casp8*^−/−^*Mlkl*^−/−^=10. Data and statistical analyses are representative of at least two experiments (A–H) or pooled from two experiments (J–M). Dashed line indicates limit of detection. ns, *p* > 0.05; \**p* < 0.05; \*\*\**p* < 0.001; \*\*\*\**p* < 0.0001; two-way ANOVA (A, D, and H), one-way ANOVA (C and F), two-tailed unpaired t test (I, J, and L), and survival curve analysis (K and M). See also Figure S9.

Given the essential role of neutrophils, we hypothesized that neutrophils could activate NLRP3 independently of caspase-8. Using bone marrow-derived neutrophils (BMNs), we demonstrated that WT neutrophils release IL-1β in response to *Fp*^CGD^ infection, which, like that of BMMs, was dependent on NLRP3 (Figures 7D, 7E, and S9O). In contrast, unlike BMMs, priming was not required for IL-1β release in neutrophils (Figures 7F and S9P). Further contrasting these cell types, pro-IL-1β expression and IL-1β release in neutrophils did not require caspase-8 (Figures 7G, 7H, and S9Q). These results suggest that neutrophils could serve as the primary in vivo source of IL- 1β in caspase-8-deficient mice.

To explore this hypothesis, we measured pro-IL-1β expression by flow cytometry in WT and *Casp8*^−/−^*Ripk3*^−/−^ mice following *Fp*^CGD^ infection. Pro-IL-1β levels were significantly reduced in macrophages and monocytes from *Casp8*^−/−^*Ripk3*^−/−^ mice compared to WT controls, whereas neutrophils showed no significant difference (Figure 7I). This supports the hypothesis that neutrophils are the primary source of IL-1β in the absence of caspase-8. However, given that baseline pro-IL-1β levels in macrophages and monocytes remain substantial in *Casp8*^−/−^*Ripk3*^−/−^ mice, we cannot exclude a residual contribution from these myeloid populations in vivo, highlighting the importance of this in vivo validation.

### An essential role for inflammasome signaling in myeloid cells during *Fp*^CGD^ infection

To test whether inflammasome signaling in myeloid cells is critical for *Fp*^CGD^ control in the absence of caspase-8 and MLKL, we generated *Casp8*^−/−^*Mlkl*^−/−^ mice with myeloid-specific deficiencies in *Asc* or *Casp1* using floxed alleles and *LysM*^Cre^-driven deletion^60,61^. *Asc*^ΔLysM^*Casp8*^−/−^*Mlkl*^−/−^ and *Casp1*^ΔLysM^*Casp8*^−/−^*Mlkl*^−/−^ mice were highly susceptible to *Fp*^CGD^ infection, displaying elevated bacterial burdens in target organs and 100% mortality (Figures 7J-7M). These phenotypes were comparable to the full-body knockout equivalent mice (Figure 1H). Similarly, myeloid-specific *Asc* deletion in *Casp8*^−/−^*Mlkl*^−/−^ mice rendered them susceptible to *Listeria monocytogenes* infection (Figure S9R). These results indicate that neutrophils, and perhaps macrophages, correctly detect the virulence capacity of *Fp*^CGD^ via NLRP3 detection, and utilize IL-1β to orchestrate an effective antibacterial immune response.

Collectively, these results establish that neutrophils are the critical cellular mediators of *Fp*^CGD^ clearance. In contrast, macrophages are largely dispensable and serve as replicative niches in susceptible hosts. Mechanistically, NLRP3 inflammasome signaling becomes essential in the absence of caspase-8 and MLKL, where it drives IL-1β production to antagonize RIPK3-driven type I interferon responses. This RIPK3-mediated interferon signaling otherwise impedes the neutrophil response (Figure S9S).

## Discussion

Regulated cell death pathways are widely viewed as essential antimicrobial defense mechanisms that eliminate intracellular replication niches. Pyroptosis, apoptosis, and necroptosis all kill cells and should prevent intracellular infection. Of these, pyroptosis and necroptosis are both lytic and thus hypothetically might be interchangeable. Strikingly, our results show the opposite—caspase- 1 and RIPK3 do not function redundantly, but instead act antagonistically by driving the release of IL-1β and type I interferon, respectively.

Like other members of the genus *Francisella*, *Fp*^CGD^ is an intracellular pathogen that replicates within myeloid cells. We initially hypothesized that regulated cell death was required to confine bacteria within pore-induced intracellular traps (PITs) created after cell death^32^, enabling their subsequent clearance through efferocytosis and killing by ROS-producing phagocytes. Initially consistent with this idea, mice lacking caspase-1 (pyroptosis), caspase-8 (apoptosis), and MLKL (necroptosis) were highly susceptible to infection. Unexpectedly, deletion of *Ripk3* completely restored resistance despite the continued absence of all three cell death pathways. Likewise, replacing *Casp8*^−/−^ with caspase-8 mutants that cannot initiate apoptosis but retain RIPK3- suppressive activity protected the mice from infection. These results demonstrate that the critical determinant of susceptibility is RIPK3-dependent signaling rather than the ability to trigger cell death. Our findings challenge the prevailing assumption that elimination of infected cellular niches is necessarily the dominant protective output of regulated cell death pathways during intracellular infection.

The central finding of this study is that the outcome of *Fp*^CGD^ infection is determined by competition between cytokines arising from two distinct cell death signaling pathways (pyroptotic and necroptotic). On one side, inflammasome activation promotes IL-1β production, neutrophil responses, and bacterial clearance. On the other, RIPK3 signaling drives type I interferon production and susceptibility. Our results support a model in which these programs are mutually antagonistic: IL-1β promotes neutrophil recruitment, whereas type I interferon suppresses it^62^. Blockade of IFNAR signaling and depletion of pDCs each protected susceptible mice, whereas induction of type I interferon rendered resistant mice susceptible. These findings establish type I interferon as a major driver of susceptibility in this setting and identify pDCs as one cellular source of this detrimental signal. The balance between these two signals therefore governs whether sufficient neutrophils are recruited and activated to achieve bacterial clearance.

Although IL-1β is essential in mice lacking caspase-8 and MLKL, IL-1β and its entire inflammasome signaling pathway are dispensable when caspase-8 is present. To explain this, we propose that basal innate signaling, likely through TLRs and TNF, provides *sufficient* activation to drive neutrophil recruitment and ROS-dependent bacterial killing. IL-1β strengthens this response and increases its robustness against perturbation^63^. Inflammasome activation therefore represents an appropriate immune response to *Fp*^CGD^ that increases the margin of host protection; however, this increased margin is only needed in the presence of an immunologic perturbation. This interpretation is supported by the observation that single deficiencies in NLRP3 pathway components result in only modest increases in bacterial burden at high challenge doses, with subsequent resolution of infection. The inflammasome pathway is activated and *appropriate*, but operates primarily as an amplifier within a larger innate defense network. However, when the immune response is perturbed by *inappropriate* type I interferon responses, the IL-1β response becomes essential to counteract the immune dysregulation, thereby restoring effective host defense.

The ability of RIPK3 to trigger the expression of type I interferons is an understudied aspect of RIPK3 signaling capacity. Increasing evidence indicates that RIPK3 can regulate inflammatory and interferon gene expression through kinase-independent scaffolding functions^64^. In our study, the type I interferon signaling capacity of RIPK3 does not require its kinase activity, as pharmacological RIPK3 kinase inhibition did not alleviate susceptibility. This suggests a requirement for the RIPK3 RHIM domain. Upstream of RIPK3, several proteins have RHIM domains and could activate RIPK3, including RIPK1, TRIF, and ZBP1. Caspase-8 deficiency is also required to trigger this signaling event. One possibility is that caspase-8 deficiency permits the accumulation of endogenous nucleic acid ligands, including Z-form nucleic acids, that engage ZBP1^65,66^. Another non-exclusive possibility is that caspase-8 normally cleaves and inactivates RIPK3^67^. The downstream pathway by which RIPK3 drives type I interferon production also remains to be elucidated.

Our model raises a conundrum because in bone marrow-derived macrophages (BMMs), caspase- 8 has previously been reported to be required for both pro-IL-1β expression and NLRP3 activation^58^, and therefore, deleting *Nlrp3* should have no effect in mice where *Casp8* was already deleted. However, our results demonstrate that susceptibility is created only when these deficiencies are combined. We sought to understand which cell type successfully activates NLRP3 in the context of a *Casp8* deletion. Myeloid-specific deletion of *Asc* or *Casp1* in *Casp8*^−/−^*Mlkl*^−/−^ mice phenocopied the susceptibility observed in complete knockouts, demonstrating that the myeloid compartment is the essential in vivo source of protective IL-1β in *Casp8*^−/−^ hosts. One interpretation is that neutrophils represent the dominant source of IL-1β in vivo, as *LysM*-Cre efficiently targets the majority of neutrophils^68^, and neutrophils retain the ability to activate NLRP3 and express pro-IL-1β independently of caspase-8. This caspase-8 independence in neutrophils may reflect their specialized transcriptional program, which enables rapid pro-IL-1β synthesis and inflammasome activation upon pathogen encounter without requiring prior TLR priming. It is therefore tempting to simply assign IL-1β release to neutrophils. However, we observed only modest reductions in pro-IL-1β expression in monocytes and macrophages in vivo, raising the possibility that the BMM phenotype in vitro does not fully reflect the responses of monocytes and macrophages in vivo. Thus, it may be that multiple myeloid populations, including resident macrophages, inflammatory monocytes, and neutrophils, contribute to the protective IL-1β response during *Fp*^CGD^ infection.

Effector-triggered immunity (ETI), originally established in plant immunity, describes host defense systems that detect the activities of pathogen virulence factors rather than the pathogens themselves^69^. In mammals, several innate immune pathways similarly respond to perturbations of core cellular processes that are commonly targeted by microbial effectors. In fact, inhibition (or deletion) of caspase-8 triggers an ETI response through RIPK3 and MLKL, resulting in necroptosis. Simultaneous deletion of caspase-8 and MLKL may also resemble the action of viral virulence factors, as herpesviruses and other viruses encode proteins that inhibit caspase -8 or suppress MLKL activation^70,71^. Our findings suggest that RIPK3 can function as an ETI sensor that responds to the disruption of caspase-8 and MLKL, initiating a compensatory type I interferon response. RIPK3 undoubtedly evolved to detect such interference by viruses. During *Fp*^CGD^ infection, however, the bacterium is unlikely to directly target caspase-8 or MLKL. Instead, the activation of this RIPK3 ETI pathway was artificially induced through genetic deletion of *Casp8* and *Mlkl* by the researchers.

Consistent with this model, the beneficial effect of RIPK3 during HSV-1 infection was independent of MLKL and could be explained by a boosted type I interferon response. The evolutionary implications are intriguing: MLKL has been independently lost in multiple vertebrate lineages, including some fish species and members of the order Carnivora, while RIPK3 has been retained^72^. Under the ETI model, loss of MLKL while retaining RIPK3 would preserve the ability of RIPK3 to detect caspase-8 inhibition and produce type I interferon while eliminating its necroptotic execution function. A speculative yet intriguing hypothesis is that the order Carnivora is more frequently exposed to diverse viruses of prey animals, and as such frequently encounters viruses that attack caspase-8 but ultimately pose a low threat level due to being mostly adapted to the prey species. A more conservative ETI response consisting of RIPK3-driven type I interferon production therefore may be less needlessly destructive than RIPK3-driven necroptosis.

These findings contribute to a growing body of evidence that type I interferons, while essential for antiviral immunity, can impair antibacterial host defenses^73,74^. Our study reveals an unexpected antagonism between the caspase-1 and RIPK3 signaling pathways. Rather than functioning redundantly to achieve cell death, caspase-1 signaling protects against bacterial infection through IL-1β-mediated suppression of a detrimental RIPK3-dependent type I interferon program. These findings establish cytokine instruction as a dominant output of these signaling pathways and suggest a broader framework in which the balance between antibacterial and antiviral immune programs governs the outcome of infection.

## Key resources and methods

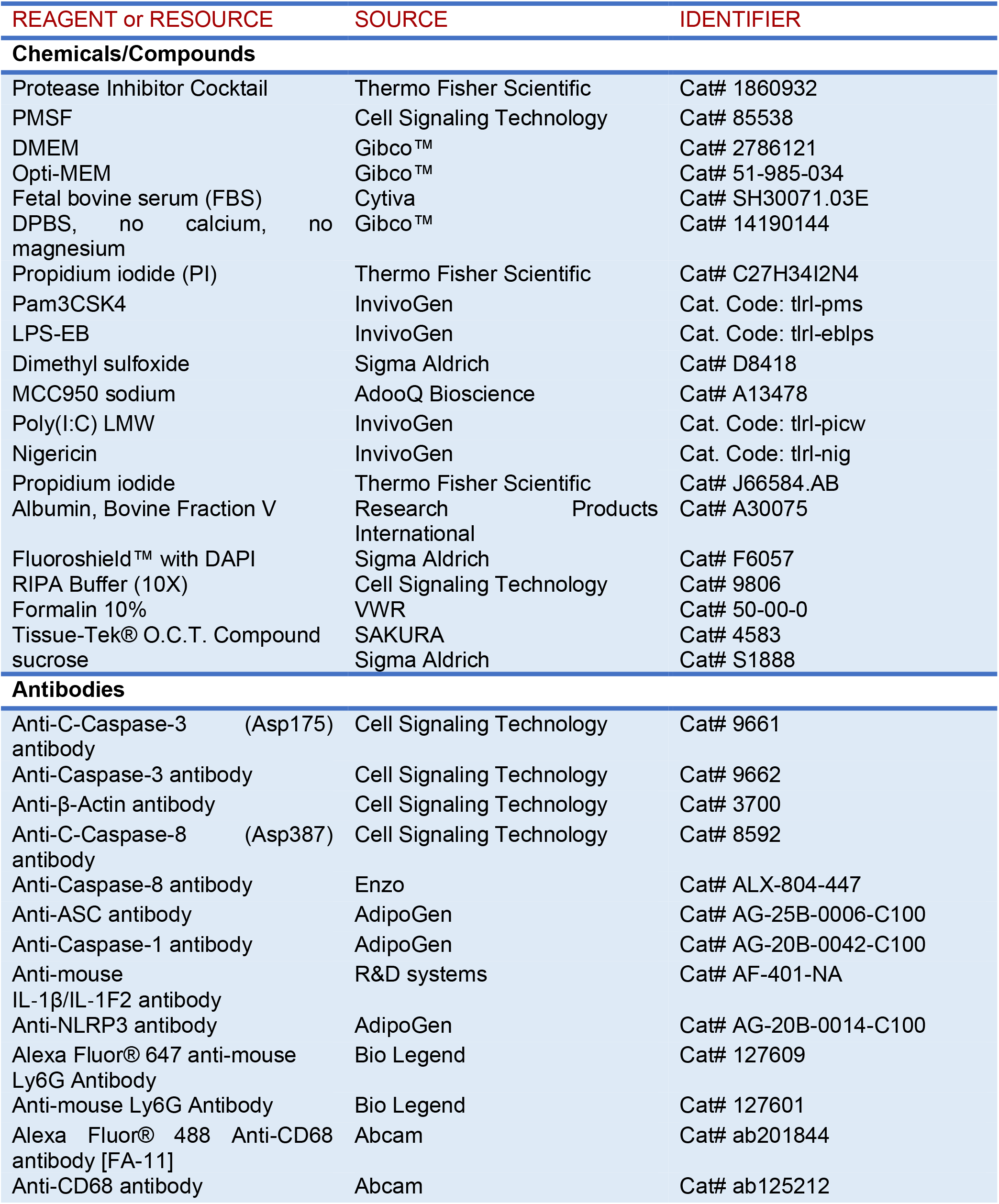
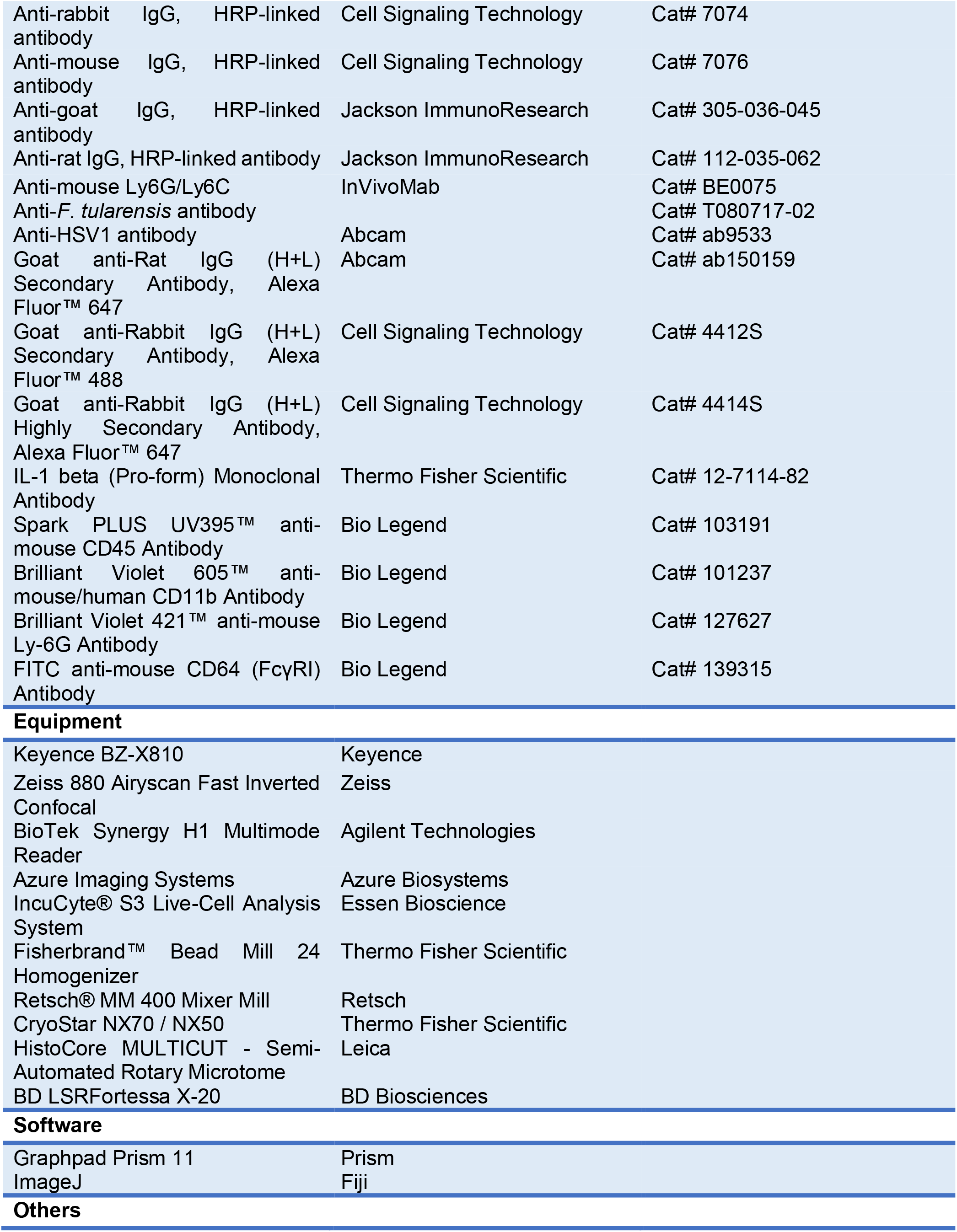
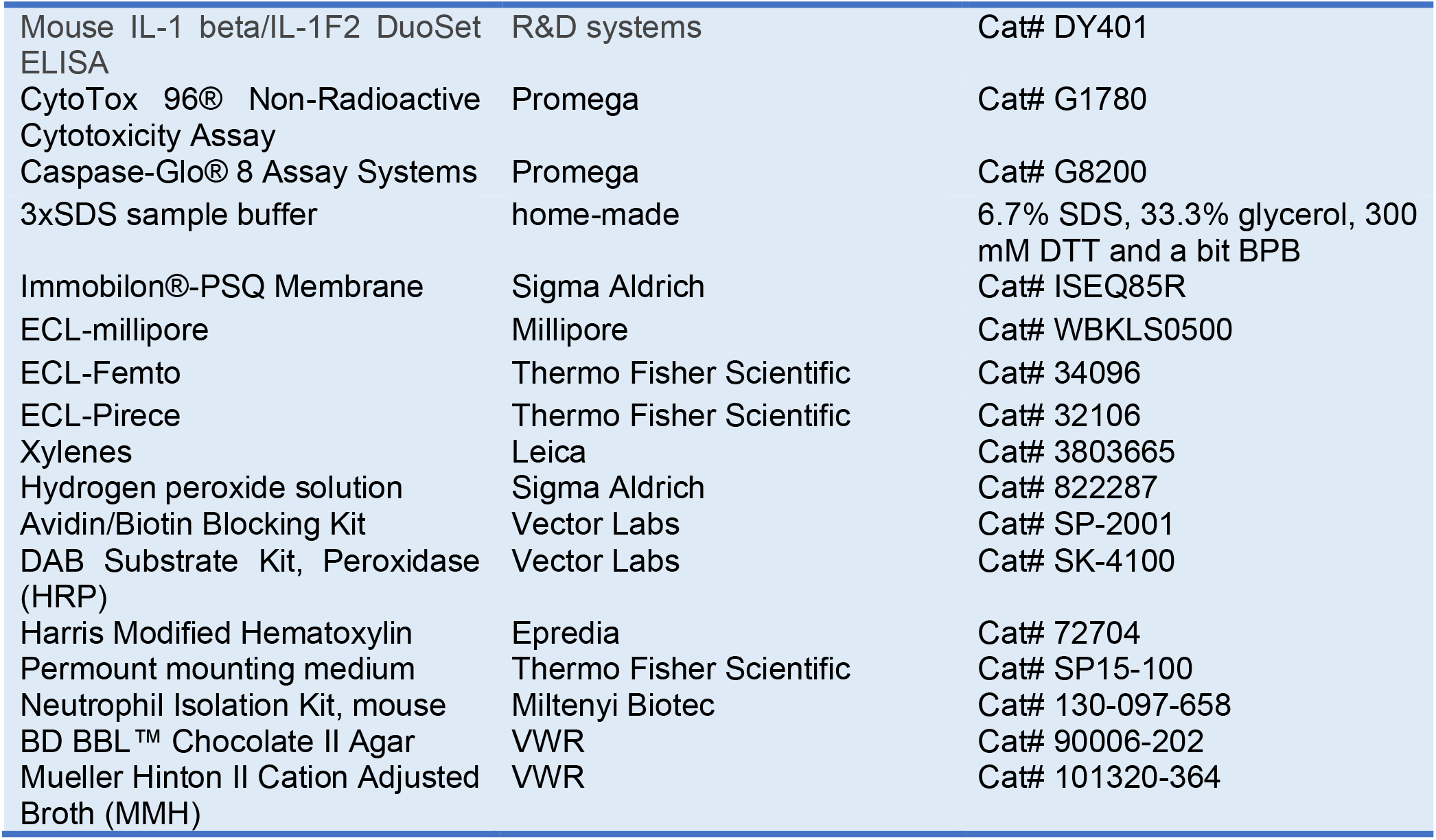

### Mouse strains

Mice were housed in a specific-pathogen-free facility at Duke University: wild-type C57BL/6 (Jackson #000664), *Rag1*^−/−^ (Jackson #002216), complement *C3*^−/−^ (Jackson #029661), complement *C7*^−/−^ (Jackson #042133), *cGAS*^−/−^ (Jackson #026554 and ref.^75^), *Ifnar*^−/−^ (Jackson #028288), *Ifng*^−/−^ (Jackson #002287), ASC-citrine (Jackson #030744), *Sting*^−/−^ (Jackson #036638), *Ifnar*^−/−^*Ifngr*^−/−^ (Jackson #029098), *Ifnlr*^−/−^ (ref.^76^), *Ncf1*^−/−^ (Jackson #004742 and ref.^77^), *Nos2*^−/−^ (Jackson #002609), *Aim2*^−/−^ (Jackson #013144), *Nlrp3*^−/−^ (ref.^78^), *Casp1*^−/−^ (Jackson #032662 and ref.^79^), *Gsdmd*^−/−^ (Jackson #32663), *Il1b*^−/−^ (ref.^80^), *Il18*^−/−^ (Jackson #004130), *Il1b*^−/−^*Il18*^−/−^ (ref.^81^), *Casp1*^fl/fl^ (ref.^60,82^), *Asc*^fl/fl^ (ref.^61^), *LysM*-Cre (Jackson #004781), *Mlkl*^−/−^ (ref.^83^), *Ripk3*^−/−^ (ref.^84^), *Ripk3*^KD^ (ref.^85^), *Ccr2*^−/−^ (ref.^22^), *Casp8*^−/−^*Mlkl*^−/−^ (ref.^86^), *Casp8*^−/−^*Ripk3*^−/−^ (ref.^46^), *Casp8*^FGLG^*Mlkl*^−/−^ (ref.^39^), *Casp8*^DA^*Mlkl*^−/−^ (ref.^39^), *Asc*^−/−^ (ref.^87^), *Casp1/11*^−/−^ (ref.^88^), *Casp7*^−/−^ (Jackson #006237). Other compound knockout mice were generated by crossing the aforementioned knockout mice in-house. Animal protocols were approved by the Institutional Animal Care and Use Committee (IACUC) at Duke University and met the US National Institutes of Health guidelines.

### Bacterial strain and growth conditions

*Francisella philomiragia* originally isolated from a CGD patient (*Fp*^CGD^) was used in this study^18^. *Fp*^CGD^ was initially streaked onto Chocolate II Agar and incubated overnight at 37°C. Single colonies were subsequently subcultured overnight at 37°C in 2 mL of Modified Mueller-Hinton (MMH) medium (composed of 21 g/L Mueller-Hinton Broth, 0.138 g/L CaCl_2_, and 0.21 g/L MgCl_2_·6H_2_O). *Francisella novicida* U112 was cultured under identical conditions. For the other bacterial strains used in this study, *Listeria monocytogenes* wild-type (strain 10403s) and the isogenic Δ*actA* (on the 10403s background) were routinely cultured in Brain Heart Infusion (BHI) broth. *Salmonella enterica* serovar Typhimurium Δ*aroA* (on the SL1344 background) was cultured in Luria-Bertani (LB) broth. All cultures were incubated at 37°C with continuous agitation. Overnight cultures were used for in vivo infections for all species.

### In vitro infections

*Fp*^CGD^ and *Fn* were grown overnight in MMH medium at 37°C with continuous agitation. Bacteria were pelleted at 12,000 × *g* for 2 min at room temperature, and the pellets were resuspended in PBS. The bacteria were then added to the cells, centrifuged at 1,000 × *g* for 10 min at room temperature, and incubated at 37°C with 5% CO_2_ for the indicated times. Gentamicin was not required because the bacteria cannot replicate extracellularly in DMEM.

### Mouse bacterial infection, survival, and CFU determination

All animal experiments used 8- to 12-week-old mice unless otherwise specified. Because no significant sex-dependent differences were observed in the infection models, both males and females were included. For *Francisella* challenges, mice were infected with the indicated colony-forming units (CFU) via intraperitoneal (IP) injection. For systemic challenges with other pathogens, mice received intravenous (IV) injections at the following target doses: 2 × 10^2^ CFU of *Salmonella* Typhimurium Δ*aroA*, 5 × 10³ CFU of wild-type *Listeria monocytogenes*, or 1 × 10⁷ CFU of *L. monocytogenes* Δ*actA*. Following inoculation, animals were monitored daily for survival and clinical signs of disease at the indicated time points. To determine bacterial burdens, mice were euthanized at specified time points post-infection. Livers and spleens were harvested aseptically, immediately transferred into 1 mL of ice-cold phosphate-buffered saline (PBS), and weighed. Organs were mechanically homogenized, and the resulting homogenates were serially diluted in cold PBS. Dilutions were plated onto appropriate selective agar media and incubated overnight at 37°C. Bacterial burdens were calculated and expressed as CFU per gram of tissue or CFU per organ.

### HSV-1 growth and infection

HSV-1 strain SC16 (kindly provided by Dr. Ashley Moseman, Duke University) was grown on Vero cells (African green monkey kidney epithelial cells; kindly provided by Dr. Raphael H. Valdivia, Duke University) as described previously^89^. Briefly, cells were cultured in DMEM supplemented with 20 mM HEPES, 10% FBS, and 100 I.U./mL penicillin-streptomycin. Viral stocks were prepared by infecting Vero cells and harvesting both the supernatant and the infected cells. Viral titers were determined via plaque assays on Vero cells. For mouse infection experiments, 8- to 12-week-old mice were infected intraperitoneally at 5 × 10^6^ PFU. Following infection, mice were monitored for survival or euthanized at designated time points for PFU determination and IHC analysis (see IHC method below). For PFU determination, liver and brain tissues were harvested post-euthanasia, minced with scissors, and homogenized using a pestle. The tissue homogenates were resuspended in DMEM medium and titrated on Vero cells.

### Bone marrow transplantation

Mice received a single dose of lethal myeloablative X-ray irradiation (900 cGy) using a CIX3 cabinet irradiator. Within 24 hours post-irradiation, mice were reconstituted via lateral tail vein injection with 1 × 10^6^ MHC haplotype-matched bone marrow cells suspended in 100 µL of sterile PBS. To prevent opportunistic infections, mice received antibiotic chow (5TK4; sulfamethoxazole and trimethoprim) from one week prior to irradiation until three weeks post-reconstitution. Following an 8-week post-transplant recovery period, mice were utilized for infection challenges.

### Bone marrow-derived macrophage isolation

Isolation of bone marrow-derived macrophages (BMMs) was performed as described previously^90^. Briefly, the two hind legs from mice were dissected post-euthanasia and disinfected with 70% ethanol. After removing all surrounding muscle tissue and fat, the ends of the cleaned bones were carefully excised to expose the marrow cavity. The bone marrow was then flushed with DMEM with 2% FBS into a collection tube using a 27-gauge needle. The resulting bone marrow suspension was passed through a 70 μm cell strainer to remove tissue and bone debris and subsequently centrifuged at 400 × *g* for 5 minutes to pellet the cells. Finally, the cell pellet was resuspended in BMM culture medium (DMEM supplemented with 20% FBS, 10% L929 conditioned medium, and Primocin) and seeded onto non-tissue culture-treated Petri dishes. Cells were cultured for 7 days to allow for differentiation. The mature BMMs were either used directly for in vitro infection assays or cryopreserved in liquid nitrogen for future use.

### ELISA

Mouse IL-1β levels in cell culture supernatants were quantified using an ELISA kit (R&D Systems, Cat. # DY401) as described previously^91^. Briefly, a high-binding 96-well plate was coated with reconstituted capture antibody (4 μg/mL) and incubated overnight at room temperature. The following day, the plate was washed four times with wash buffer (0.05% Tween-20 in PBS, pH 7.2–7.4), and residual buffer was removed by inversion and blotting onto clean paper towels. Wells were blocked with blocking buffer (1% BSA in PBS, 0.22 μm filtered, pH 7.2–7.4) for 1 hour at room temperature. Recombinant mouse IL-1β standard was reconstituted in blocking buffer, and 2-fold serial dilutions were prepared starting at 1000 pg/mL. Standards and cell culture supernatants were added to the designated wells and incubated for 2 hours at room temperature. After four washes, 250 ng/mL of reconstituted detection antibody was added to each well, followed by a 2-hour incubation at room temperature. The plate was washed four times, and streptavidin-HRP diluted in blocking buffer was added to each well and incubated in the dark for 20 minutes. Following a final wash cycle, substrate solution was added to each well, and the color development was closely monitored. The reaction was terminated with stop solution once the desired color intensity was achieved. Absorbance was measured at 450 nm using a microplate reader. IL-1β concentrations were calculated using a standard curve.

### Histology

For paraffin-embedded tissues, samples were collected from mice post-euthanasia and placed in 10% formalin within 50 mL conical tubes. The tubes were rotated for at least three days to ensure complete formalin penetration into the tissue. Once fixed, the tissues were transferred to tissue cassettes for embedding, sectioning, slide mounting, and staining. For frozen tissue samples, tissues were harvested at designated time points post-infection and fixed in 2% paraformaldehyde (PFA) for 24 hours at 4°C. The organs were then equilibrated in 30% sucrose for two days and subsequently incubated in OCT compound at 4°C for 4 hours. Finally, the tissues were embedded in frozen tissue molds filled with OCT compound, frozen on dry ice, and stored at -80°C. The frozen tissues were sectioned using a cryostat.

### H&E staining

Paraffin-embedded tissue sections were baked at 55°C for 1 hour prior to staining. The slides were then deparaffinized in xylene twice (2 minutes each) with gentle agitation and rehydrated through a graded ethanol series: 100% ethanol once (1 minute), 95% ethanol once (1 minute), 80% ethanol once (1 minute), and 70% ethanol once (1 minute). To complete rehydration, the slides were immersed in PBS for 5 minutes. Sections were then stained with hematoxylin for 5 minutes and rinsed thoroughly under running tap water. The slides were immersed in bluing reagent (0.2% ammonia in deionized water) for 10–15 seconds and rinsed well with water to normalize the pH and ensure optimal eosin staining. Next, the slides were dipped in 100% ethanol and counterstained with Eosin Y for 3 minutes. Afterwards, the slides were rinsed in tap water until the eosin ceased streaking. Finally, the sections were dehydrated through a graded series of 50% ethanol (30 seconds), 70% ethanol (30 seconds), 95% ethanol (30 seconds), and 100% ethanol (30 seconds), cleared in xylene twice (1–2 minutes each), and mounted with Permount mounting medium under a glass coverslip. Images were captured using a KEYENCE BZ-X810 fluorescence microscope.

### Immunohistochemistry

For immunohistochemical analysis of paraffin-embedded tissues, 5 μm sections were deparaffinized in xylene three times (5 minutes each) and then rehydrated through a graded ethanol series: 100% ethanol three times (3 minutes each), 95% ethanol twice (3 minutes each), and 80% ethanol once (3 minutes). The slides were then immersed in double-distilled water (ddH_2_O) for 5 minutes to complete rehydration. For antigen retrieval, slides were incubated in sodium citrate buffer (2.94 g Tri-Sodium Citrate, ddH_2_O 1 L, mix to dissolve, pH to 6.0 with 1 N HCl, add 0.05 mL Tween-20) and heated in a pressure cooker (Instant Pot Ultra) on the high-pressure setting for 12 minutes. Slides were cooled in a slide rack at room temperature for a minimum of 20 minutes and then washed in 1× TBST (100 mL 10× TBS, 900 mL ddH_2_O, 1 mL Tween-20) for 1 minute. Residual liquid was removed by tapping the slides on paper towels and wiping around the tissues with Kimwipes. A hydrophobic barrier was created around the tissue sections using a PAP pen, and the specimens were blocked with normal goat serum blocking solution (2.5%; Vector Laboratories, Cat. # S-1012-50) for 30 minutes at room temperature. Following two washes with 1× TBST, the slides were incubated overnight at 4°C with primary antibodies in a humidified chamber. The next day, slides were washed three times in 1× TBST (1 minute each) and treated with 3% hydrogen peroxide for 10 minutes to block endogenous peroxidase activity. After three additional 1-minute washes, sections were incubated with an HRP-conjugated secondary antibody polymer system (SignalStain Boost IHC Detection Reagent, HRP anti-rabbit, Cell Signaling Technology, Cat. # 8114; or ImmPRESS HRP Goat Anti-Rat IgG Polymer Detection Kit, Vector Laboratories, Cat. # MP-7404-50). The slides were washed three times (5 minutes each), incubated with a DAB Substrate Kit, and the reaction was stopped by transferring the slides to ddH_2_O. Following three 1-minute washes and a 5-minute rinse in running ddH_2_O, sections were counterstained with Harris Modified Hematoxylin for 1 minute. The hematoxylin was blued under running tap water for 1 minute. Finally, the slides were dehydrated through a graded series of 50% ethanol (2 minutes), 70% ethanol (2 minutes), and 100% ethanol (twice, 2 minutes each), cleared in xylene (twice, 5 minutes each), and mounted with Permount mounting medium under a glass coverslip. Images were captured using a KEYENCE BZ-X810 microscope.

### Immunofluorescence and confocal analyses

For immunofluorescence analysis of cultured cells, BMMs were seeded onto glass coverslips. Cells were then infected with *Fp*^CGD^ or *Fn* at an MOI of 5 for the indicated time points. Following incubation, the culture medium was removed, and cells were washed three times with cold PBS. Cells were fixed with 4% PFA for 15 minutes at 4°C, followed by three additional washes with PBS. To block non-specific binding and permeabilize the samples, both frozen tissue slides and cell culture samples were incubated in a blocking/permeabilization solution (0.3% Triton X-100 and 5% goat serum in PBS) for 40 minutes at room temperature under rotation. After three washes with PBS, primary antibodies diluted in antibody dilution buffer (1% BSA and 0.3% Triton X-100 in PBS) were applied overnight at 4°C with gentle rotation: anti-cleaved caspase-3 (1:400), anti- *Ft* LPS (1:1000, which cross-reacts with *Fp*^CGD^), and anti-Ly6G (1:200). The following day, samples were washed three times with PBS (5 minutes per wash) and incubated with appropriate secondary antibodies. Finally, the samples were washed three times with PBS and mounted using Fluoroshield™ with DAPI for nuclear counterstaining. Images were captured using a Zeiss LSM 880 Airyscan Fast inverted confocal microscope or a Keyence BZ-X810 fluorescence microscope.

### Immunoblot analyses

Following *Fp*^CGD^ or *Fn* infection of BMMs for the indicated time points, the culture medium was collected. The cells were then washed with ice-cold PBS and lysed in RIPA buffer supplemented with protease inhibitors and PMSF at 4°C for 10 minutes. The lysates were centrifuged at 15,000 rpm (21,130 × *g*) for 10 minutes at 4°C to remove insoluble debris. Subsequently, 4× homemade loading buffer was added to both the cell lysates and culture medium samples. Equal amounts of protein lysates and supernatant samples were resolved on 15% SDS-PAGE gels using SDS-PAGE running buffer at a constant voltage of 128 V for 65 minutes. Separated proteins were transferred onto PVDF membranes using a Bio-Rad transfer system at 300 mA for 2 hours at 4°C. The PVDF membranes were blocked with 5% BSA in TBST for 30 minutes at room temperature and incubated with primary antibodies overnight at 4°C with gentle shaking. The following day, membranes were washed three times with TBST (10 minutes per wash) and incubated with HRP-conjugated secondary antibodies diluted in 5% BSA in TBST for 1 hour at room temperature with shaking. After three additional 10-minute washes in TBST, the membranes were incubated with ECL reagents for 5 minutes at room temperature prior to visualization using an Azure Imaging System.

### LDH assay

BMMs were seeded in 96-well plates and primed overnight with either Pam3CSK4 or LPS. On the second day, the cells were infected. At the indicated time points, the culture supernatant was collected using a multichannel pipette and stored at -80°C for subsequent assays. LDH release was measured using the CytoTox 96® Non-Radioactive Cytotoxicity Assay kit (Promega) according to the manufacturer’s instructions. Briefly, 50 μL of reconstituted CytoTox 96® substrate mix was added to each well of a new 96-well plate, followed by the transfer of 50 μL of the collected supernatant samples. The plate was incubated at 37°C in the dark while color development was monitored. The enzymatic reactions were terminated by adding stop solution. After ensuring the removal of any macroscopic bubbles, the absorbance was measured at 490 nm using a BioTek Synergy H1 Multimode Reader.

### IncuCyte cell assay

For IncuCyte live-cell imaging, BMMs were seeded at a density of 5 × 10^4^ cells per well into Corning® Falcon® cell culture plates (Corning, Cat. # CLS353072) compatible with the IncuCyte® S3 Live-Cell Analysis System and primed overnight with either Pam3CSK4 or LPS. On the following day, propidium iodide (PI) reagent was added to the wells, and the cells were infected with *Fp*^CGD^ or *Fn* at an MOI of 5. The plate was immediately placed into the IncuCyte® system for up to 24 hours. Phase-contrast and fluorescence images were captured every 30 minutes or 1 hour using a 10× objective. The fluorescence area was subsequently quantified using the IncuCyte® Cell-by-Cell Analysis Software module.

### Flow cytometry

At the indicated time points post-infection, mice were euthanized, and livers were harvested for flow cytometry. Livers were minced on ice using scissors and incubated in digestion buffer (1 U/mL Liberase TM, 0.2 mg/mL DNase I, and 50 µg/mL gentamicin in DMEM supplemented with 1 mM CaCl_2_ and 1 mM MgCl_2_) for 40 minutes in an incubation shaker with 5% CO_2_. Digested livers were homogenized through a 70 µm cell strainer, followed by washing with DMEM and centrifugation at 300 × *g* for 10 minutes at 4 °C. Samples were counted using a Countess automated cell counter according to the manufacturer’s protocol, and 5 × 10^6^ cells per tissue per mouse were stained with the indicated Fc block, antibodies, and viability dyes for 30 minutes. For pro-IL-1β, staining was performed using an Intracellular Fixation & Permeabilization Buffer according to the product manual. For each sample, 100,000 events were acquired on a BD LSR Fortessa X-20 Cell Analyzer at the Duke Flow Cytometry Core Facility. Samples were analyzed using FlowJo™ (version 10.7.1). Myeloid populations were sequentially gated from live, single CD45⁺ cells. Within the CD11b⁺ population, neutrophils were identified as Ly6G⁺CD64⁻. The Ly6G⁻ population was further resolved into CD11b^hi^CD64^lo^ monocytes and CD11b^lo^CD64^hi^ macrophages.

### Bulk RNA sequencing and analysis

Mouse tissues were harvested, snap-frozen, and homogenized in TRIzol reagent for total RNA extraction. RNA samples were submitted to Admera Health for library preparation and sequencing. RNA quality was assessed using an Agilent High Sensitivity RNA TapeStation, and RNA concentration was determined using the AccuBlue Broad Range RNA Quantitation Assay (Biotium). Poly(A)+ RNA was isolated using the NEBNext Poly(A) mRNA Magnetic Isolation Module (New England Biolabs). Libraries were prepared using the NEBNext Ultra II Directional RNA Library Prep Kit for Illumina (New England Biolabs) according to the manufacturer’s protocol. Library quality was evaluated by Agilent TapeStation D1000 ScreenTape and quantified using a Qubit 2.0 fluorometer (Thermo Fisher Scientific). Libraries (average insert size ∼300 bp; final library size ∼430 bp) were dual-indexed, pooled at equimolar concentrations, and sequenced on an Illumina NovaSeq X Plus platform to generate 150-bp paired-end reads at a depth of approximately 40 million read pairs per sample.

Raw sequencing reads were assessed using FastQC (v0.12.1) and trimmed with Trimmomatic (v0.39) using default parameters. Reads were aligned to the mouse reference genome (GRCm38/mm10) using STAR (v2.7.10b). PCR duplicates were marked with Picard (v3.0.0), transcript assembly was performed using StringTie (v2.2.1), and gene-level read counts were generated with HTSeq (v2.0.3).

Differential gene expression analysis was performed using RNA-seq Explorer v37 (https://zoeliu-duke.github.io/RNAseqExplorer/). Initial quality control and exploratory analyses were conducted using a browser-based interface. For final statistical analysis, count matrices were exported and analyzed locally in R using the Bioconductor package DESeq2. Differential expression statistics and variance-stabilized expression values were then imported back into RNA-seq Explorer for visualization, pathway enrichment, and transcription factor activity analyses.

### Caspase-8 activity assay

BMMs were seeded at a density of 5 × 10^4^ cells per well into white-walled 96-well plates. On the following day, the cells were infected with *Fp*^CGD^ at an MOI of 5 for the indicated time points. Prior to the assay, the plates were removed from the incubator and allowed to equilibrate to room temperature. Next, 100 μL of Caspase-Glo® 8 Reagent was added to each well containing 100 μL of blank, negative control, or infected cell culture medium. The plates were covered with lids, gently mixed on a horizontal plate shaker at 500 rpm for 2 minutes, and then incubated at room temperature for 3 hours in the dark. Finally, luminescence was measured using a BioTek Synergy H1 Multimode Reader.

### Bone marrow-derived neutrophil isolation

Isolation of bone marrow-derived neutrophils (BMNs) was performed as described previously^79^. Briefly, hind legs were dissected post-euthanasia and disinfected with 70% ethanol. After removing all surrounding muscle and adipose tissue, the ends of the cleaned bones were carefully excised to expose the marrow cavity. The bone marrow was then flushed with DMEM into a collection tube using a 27-gauge needle. The resulting bone marrow suspension was passed through a 40 μm cell strainer to remove tissue and bone debris, and total cell counts were determined.

Neutrophil isolation was performed using a commercial Neutrophil Isolation Kit according to the manufacturer’s instructions. Briefly, the cell suspension was centrifuged at 300 × *g* for 10 minutes, and the supernatant was discarded. The cell pellet was resuspended in ice-cold isolation buffer (PBS, pH 7.2, 0.5% BSA, and 2 mM EDTA) at a ratio of 200 μL per 5 × 10^7^ cells. A neutrophil biotin-antibody cocktail (50 μL per 5 × 10^7^ cells) was added, and the mixture was incubated on ice for 10 minutes. Cells were then washed with 10 mL of buffer per 5 × 10^7^ cells, centrifuged at 300 × *g* for 10 minutes, and resuspended in 400 μL of buffer per 5 × 10^7^ cells. Afterwards, anti-biotin MicroBeads (100 μL per 5 × 10^7^ cells) were added, followed by a 15-minute incubation on ice. Cells were washed again with 10 mL of buffer per 5 × 10^7^ cells, centrifuged, and resuspended in 500 μL of buffer for up to 1 × 10^8^ cells.

For the magnetic separation step, the provided column was placed in the magnetic field of a MACS separator. The cell suspension was applied to the column, and the flow-through— containing unlabeled, enriched neutrophils—was collected. The column was washed with buffer, and the flow-through was pooled. Total cell numbers in the collected fraction were determined before the suspension was centrifuged at 300 × *g* for 10 minutes. Finally, the enriched BMNs were resuspended in OPTI-MEM medium and seeded into 96-well or 6-well plates for subsequent *Fp*^CGD^ infection assays.

## Statistics

All statistical analyses were performed using GraphPad Prism (version 11.0.2). Statistical significance was determined using an unpaired two-tailed Student’s t test for comparisons between two groups, one-way ANOVA with Tukey’s multiple-comparisons test for comparisons among three or more groups, two-way ANOVA with Tukey’s multiple-comparisons test for experiments involving two independent variables, or the log-rank (Mantel–Cox) test for survival analyses. Statistical significance was defined as follows: ns, *p* > 0.05; \**p* < 0.05; \*\**p* < 0.01; \*\*\**p* < 0.001; \*\*\*\**p* < 0.0001.

## Acknowledgements

We thank E. Ashley Moseman, Nicholas S. Heaton, Francis Ka-Ming Chan, Mari L. Shinohara, Pengda Liu, Richard A. Flavell, and Xiaoxia Li for sharing mice. Flow cytometry was performed in the Duke Cancer Institute Flow Cytometry Facility at Duke University, Durham, NC, which is supported by the NCI Cancer Center Support Grant (CCSG) award number P30CA014236. The authors gratefully acknowledge the Duke Light Microscopy Core Facility and the Duke Small Animal Irradiator Service Center for their support and assistance in this work. This work was supported by NIH grants AI136920, AI175078, AI181815, and AI136920 (E.A.M.).

---

**Figure S1. Several immune pathways are dispensable for resistance to *Fp*^CGD^ infection.** Mice were infected intraperitoneally (IP) with 10^5^ CFU of *Fp*^CGD^. CFU burdens in the liver and spleen were determined at the indicated time points post-infection (A, B, E, F, and H). Survival was monitored (C, D, G, I, and J). Mice: (A) n=4 for all; (B) WT=4, *Rag1*^−/−^=3; (C) WT=5, *Rag1*^−/−^=3, *Ncf1*^−/−^=3; (D) WT=5, Complement *C3*^−/−^=3; (E) WT=4, Complement *C7*^−/−^=3; (F) n=5 for all; (G) WT=5, *Sting*^−/−^=3, *Ifnar*^−/−^=3; (H) WT=11, *Ifng*^−/−^=9, *Ifnar*^−/−^*Ifngr*^−/−^=12; (I) n=5 for both; (J) WT=10, *Ifng*^−/−^=5, *Ifnar*^−/−^*Ifngr*^−/−^*Ifnlr*^−/−^=5. Data and statistical analyses are representative of at least two experiments (B, G, and I), pooled from two experiments (C, F, and J), or pooled from three experiments (H). Dashed line indicates limit of detection. ns, *p* > 0.05; \**p* < 0.05; \*\**p* < 0.01; \*\*\**p* < 0.001; two-tailed unpaired t test (B and E), survival curve analysis (C, D, G, I, and J), and one-way ANOVA (F and H).

**Figure S2. Characterization of NLRP3 inflammasome activation by *Fp*^CGD^.** Bone marrow-derived macrophages (BMMs) were left unprimed or primed overnight with Pam3CSK4 (500 ng/mL) or LPS (100 ng/mL) and subsequently infected with *Fp*^CGD^ or *Fn* at an MOI of 5 for the indicated time points. IL-1β release was measured by ELISA at the indicated time points (A, B, C, F, J, L, O, Q, and R). Western blotting was performed at 4 HPI (E and N). PI uptake was measured using IncuCyte (D, H, and I). Cell death was quantified by LDH release (G, K, M, and P). Live or heat-killed (HK; 65°C for 10 min) *Fp*^CGD^ was used in (O and P). BMMs were pretreated for 30 min with KCl (130 mM) (Q) or NAC (5 mM) (R) prior to infection with *Fp*^CGD^ at an MOI of 5. Data and statistical analyses are representative of at least two experiments. ns, *p* > 0.05; \*\*\**p* < 0.001; \*\*\*\**p* < 0.0001; two-way ANOVA (A–C, F, G, and J–M) and one-way ANOVA (D, H, I, and O–R).

**Figure S3. Inflammasome knockout mice resist *Fp*^CGD^.** Mice were infected IP with 10^5^ CFU of *Fp*^CGD^ unless otherwise indicated. CFU burdens in the liver and spleen were determined at the indicated time points post-infection (A–E, G, H, and J–M). Survival was monitored (F and I). Mice were treated daily with vehicle or MCC950 (50 mg/kg, IP) beginning on the day of infection (M). Mice: (A) n=5 for all; (B) n=4 for all; (C) WT=6, *Asc*^−/−^*Casp1/11*^−/−^=13; (D) WT=5, *Casp1/11*^−/−^*Casp7*^−/−^=4; (E) WT=7, *Gsdmd*^−/−^=8; (F) n=5 for both; (G) WT=6, *Il1b*^−/−^*Il18*^−/−^=8; (H) WT=4, *Il1b*^−/−^=4, *Il18*^−/−^=3, *Il1b*^−/−^*Il18*^−/−^=4; (I) WT=6, *Il1b*^−/−^=5, *Il18*^−/−^=5, *Il1b*^−/−^*Il18*^−/−^=6; (J) n=5 for both; (K) WT=10, *Asc*^−/−^=8; (L) WT=10, *Nlrp3*^−/−^=11; (M) DMSO=4, MCC950=5. Data and statistical analyses are pooled from two experiments (A, C–G, and I–M), or representative of at least two experiments (B and H). Dashed line indicates limit of detection. ns, *p* > 0.05; \**p* < 0.05; \*\**p* < 0.01; one-way ANOVA (A, B, and H), two-tailed unpaired t test (C–E, G, and J–M), survival curve analysis (F and I).

**Figure S4. Loss of caspase-8 and necroptosis alone does not cause susceptibility to *Fp*^CGD^ infection.** (A–M) Mice were infected IP with 10^5^ CFU of *Fp*^CGD^ unless otherwise indicated. CFU burdens in the liver and spleen were determined at the indicated time points post-infection (A–K). Survival was monitored (L). Mice: (A) WT=5, *Ripk3*^−/−^=4; (B) *Mlkl*^+/−^=5, *Mlkl*^−/−^=8; (C) n=5 for both; (D) WT=4, *Ripk3*^KD^=5, *Ncf1*^−/−^=3; (E) WT=9, *Ripk3*^KD^=6; (F) *Casp8*^+/−^*Ripk3*^−/−^=4, *Casp8*^−/−^*Ripk3*^−/−^=5; (G) *Casp8*^+/−^*Ripk3*^−/−^=6, *Casp8*^−/−^*Ripk3*^−/−^=7; (H) WT=5, *Casp8*^+/−^*Mlkl*^−/−^=8, *Casp8*^−/−^*Mlkl*^−/−^=6; (I) *Casp8*^+/−^*Mlkl*^−/−^=12, *Casp8*^−/−^*Mlkl*^−/−^=5; (J) WT=5, *Casp8*^+^*Ripk3*^−/−^=10, *Casp8*^−/−^*Ripk3*^−/−^=6, *Casp8*^+^ denotes both *Casp8*^+/+^ and *Casp8*^+/−^; (K) WT=5, *Casp8*^+/−^*Mlkl*^−/−^=9, *Casp8*^−/−^*Mlkl*^−/−^=11; (L and M) n=5 for all. (N) BMMs were left unprimed or primed overnight with Pam3CSK4 (500 ng/mL) and subsequently infected with *Fp*^CGD^ at an MOI of 5. Cell lysates were collected at 4 HPI and analyzed by western blotting. Data and statistical analyses are representative of at least two experiments (A–C, and N), or pooled from two experiments (E–M). Dashed line indicates limit of detection. ns, *p* > 0.05; \*\*\*\**p* < 0.0001; two-tailed unpaired t test (A– C, E–G, and I), one-way ANOVA (D, H, J, K, and M), survival curve analysis (L).

**Figure S5. RIPK3-mediated susceptibility is not explained by lymphoproliferative disease.** Mice were infected IP with 10^5^ CFU of *Fp*^CGD^. CFU burdens in the liver and spleen were determined at the indicated time points (A, C, and I–L). (B) Spleen weights were measured. Survival was monitored (D–H). Mice were treated daily with vehicle or MCC950 (50 mg/kg, IP) beginning on the day of infection (L). Mice: (A) n=6 for both; (B) *Asc*^−/−^*Casp1/11*^−/−^*Casp8*^+/−^*Ripk3*^−/−^=6, *Asc*^−/−^*Casp1/11*^−/−^*Casp8*^−/−^*Ripk3*^−/−^=5, *Asc*^−/−^*Casp1/11*^−/−^*Casp8*^−/−^*Mlkl*^−/−^=4; (C) n=5 for both; (D) *Asc*^−/−^*Casp8*^+/−^*Ripk3*^−/−^=4, *Asc*^−/−^*Casp8*^−/−^*Ripk3*^−/−^=4, *Casp1*^−/−^*Casp8*^+/−^*Ripk3*^−/−^=4, *Casp1*^−/−^*Casp8*^−/−^*Ripk3*^−/−^=3; (E) *Asc*^−/−^*Casp1/11*^−/−^*Casp8*^+/−^*Ripk3*^−/−^=6, *Asc*^−/−^*Casp1/11*^−/−^*Casp8*^−/−^*Ripk3*^−/−^=11; (F) WT=5, *Asc*^−/−^*Casp8*^+/−^*Mlkl*^−/−^=8, *Asc*^−/−^*Casp8*^−/−^*Mlkl*^−/−^=6; (G) *Asc*^−/−^*Casp1/11*^−/−^*Casp8*^+/−^*Mlkl*^−/−^=14, *Asc*^−/−^*Casp1/11*^−/−^*Casp8*^−/−^*Mlkl*^−/−^=6; (H) n=6 for both; (I) WT=5, *Asc*^−/−^*Casp8*^+/−^*Ripk3*^−/−^*Mlkl*^−/−^=5, *Asc*^−/−^*Casp8*^−/−^*Ripk3*^−/−^*Mlkl*^−/−^=6; (J) WT=5, *Asc*^+/−^*Casp8*^FGLG^*Mlkl*^−/−^=7, *Asc*^−/−^*Casp8*^FGLG^*Mlkl*^−/−^=12; (K) WT=5, *Asc*^−/−^*Casp8*^DA^*Mlkl*^−/−^=4; (L) DMSO=4, MCC950=5. Data and statistical analyses are pooled from two experiments (A–J, and L) or representative of two experiments (K). Dashed line indicates limit of detection. ns, *p* > 0.05; \*\**p* < 0.01; \*\*\**p* < 0.001; \*\*\*\**p* < 0.0001; two-tailed unpaired t test (A, C, K, and L), survival curve analysis (D, and E–H), and one-way ANOVA (B, I, and J).

**Figure S6. Differential requirements for cell death pathways across bacterial infection models.** Mice were infected intravenously with *Lm* (5 × 10^3^ CFU; A–F), or *Lm* Δ*actA* (1 × 10^7^ CFU; G). Survival was monitored (A, C, and D). CFU burdens in the liver and spleen were determined at the indicated time points post-infection (B and E–G). Mice: (A) WT=5, *Asc*^−/−^*Casp1/11*^−/−^*Casp8*^+/−^*Mlkl*^−/−^=4, *Asc*^−/−^*Casp1/11*^−/−^*Casp8*^−/−^*Mlkl*^−/−^=3; (B) *Casp1*^−/−^*Casp8*^+^*Mlkl*^−/−^=7, *Casp1*^−/−^*Casp8*^−/−^*Mlkl*^−/−^=4, *Casp8*^+^ denotes both *Casp8*^+/+^ and *Casp8*^+/−^; (C) WT=7, *Casp1*^−/−^*Casp8*^+/−^*Mlkl*^−/−^=10, *Casp1*^−/−^*Casp8*^−/−^*Mlkl*^−/−^=7; (D) *Casp1*^−/−^*Casp8*^+/−^*Ripk3*^−/−^=5, *Casp1*^−/−^*Casp8*^−/−^*Ripk3*^−/−^=9; (E) WT=10, *Casp8*^+/−^*Ripk3*^−/−^=6, *Asc*^−/−^*Casp8*^+/−^*Ripk3*^−/−^=12, *Casp8*^−/−^*Ripk3*^−/−^=6, *Asc*^−/−^*Casp8*^−/−^*Ripk3*^−/−^=13; (F) *Asc*^−/−^*Casp8*^+/−^*Mlkl*^−/−^=5, *Asc*^−/−^*Casp8*^−/−^*Mlkl*^−/−^=4; (G) WT=5, *Asc*^−/−^*Casp8*^+/−^*Ripk3*^−/−^=6, *Asc*^−/−^*Casp8*^−/−^*Ripk3*^−/−^=4, *Asc*^−/−^*Casp8*^+/−^*Mlkl*^−/−^=5, *Asc*^−/−^*Casp8*^−/−^*Mlkl*^−/−^=5. Data and statistical analyses are pooled from two experiments. Dashed line indicates limit of detection. ns, *p* > 0.05; \**p* < 0.05; \*\**p* < 0.01; \*\*\**p* < 0.001; \*\*\*\**p* < 0.0001; survival curve analysis (A, C, and D), two-tailed unpaired t test (B and F), and one-way ANOVA (E and G).

**Figure S7. RIPK3 kinase activity is dispensable, whereas plasmacytoid dendritic cells contribute to susceptibility.** (A) Mice were treated daily with an isotype control antibody or a combination of anti-IFNAR and anti-IFNGR antibodies (0.2 mg/antibody/dose, IP) beginning at −1 DPI, infected IP with 10^7^ CFU of *Fp*^CGD^, and CFU burdens were determined at 3 DPI. Mice: n=8 for both. (B) Mice were untreated or treated daily with vehicle or GSK-872 (8 mg/kg, IP) beginning on the day of infection, infected IP with 10^5^ CFU of *Fp*^CGD^, and CFU burdens were determined at 5 DPI. Mice: *Asc*^−/−^*Casp1/11*^−/−^*Casp8*^+/−^*Mlkl*^−/−^=12, *Asc*^−/−^*Casp1/11*^−/−^*Casp8*^−/−^*Mlkl*^−/−^+Vec=9, *Asc*^−/−^*Casp1/11*^−/−^*Casp8*^−/−^*Mlkl*^−/−^+GSK-872=9. (C) Mice were treated daily with an isotype control antibody or anti-BST2 antibody (0.2 mg/dose, IP) beginning at −1 DPI, infected IP with 10^5^ CFU of *Fp*^CGD^, and CFU burdens were determined at 4 DPI. Mice: n=6 for both. (D) Mice were infected IP with HSV-1 strain SC16 (5 × 10^6^ PFU). Liver and brain sections were collected at 5 DPI and analyzed by IHC for HSV-1; images shown are representative from mice with significant symptomatic infection. Mice: n=4 for both. Data and statistical analyses are pooled from two experiments. Dashed line indicates limit of detection. ns, *p* > 0.05; \**p* < 0.05; \*\**p* < 0.01; two-tailed unpaired t test (A and C) and one-way ANOVA (B). Scale bars: 50 μm in (D).

**Figure S8. Supporting evidence for neutrophil-dependent protection against *Fp*^CGD^.** (A and B) Mice were infected IP with 10^5^ CFU of *Fp*^CGD^. CFU burdens in the liver and spleen were determined at 2 DPI (A), and survival was monitored (B). Mice: (A) n=4 for both; (B) n=5 for both. (C) Mice were treated with control liposomes or clodronate liposomes (200 μL, IV, every other day) beginning at −1 DPI, infected IP with 10^5^ CFU of *Fp*^CGD^, and CFU burdens were determined at 4 DPI. Mice: n=4 for both. (D–G) Mice were infected IP with 10^7^ CFU of *Fp*^CGD^. Livers were collected at 8 HPI for immunofluorescence staining of *Fp*^CGD^ (red) (D). Mice received an isotype control antibody or an anti-Ly6G antibody (0.2 mg/dose, IP, daily) beginning at −1 DPI. Spleens were collected at 1 DPI for H&E staining and IHC staining of Ly6G, CD68, and *Fp*^CGD^ (E). Spleens from naïve mice, as well as infected mice at 12 HPI and 5 DPI, were snap-frozen for RNA extraction using TRIzol, followed by bulk RNA sequencing. The top 50 differentially expressed genes are shown (F). Mice: (D and E) n=3; (F) n=3–6 for each group; (G) n=4 for all. (H and I) Mice received either an isotype control antibody or an anti-IFNAR antibody (0.2 mg/dose, IP, daily) beginning at −1 DPI. Mice were infected IP with 10^5^ CFU of *Fp*^CGD^. Livers (H) and spleens (I) were collected at 5 DPI for H&E staining and IHC staining of Ly6G, CD68, and *Fp*^CGD^. Mice: n=4 for all. Dashed line indicates limit of detection. ns, *p* > 0.05; \**p* < 0.05; two-tailed unpaired t test (A and C), and survival curve analysis (B). Scale bars: 25 μm in (D); 50 μm in (E) and (G–I).

**Figure S9. NLRP3 activation in bone marrow-derived neutrophils is caspase-8-independent.** (A–N) BMMs were left unprimed or primed overnight with Pam3CSK4 (500 ng/mL) or LPS (100 ng/mL) and infected with *Fp*^CGD^ or *Fn* at an MOI of 5. Cell death was quantified by LDH release (A, D, G, I, and M). IL-1β release was measured by ELISA (B, C, F, H, and L). Inflammasome activation was assessed by western blotting (E, J, and K). PI uptake was measured using IncuCyte (N). (O–Q) Bone marrow-derived neutrophils (BMNs) were infected with *Fp*^CGD^ at an MOI of 20 (O and P) or 10 (Q). Cell culture supernatants were collected at the indicated time points, and LDH release was measured. (R) Mice were infected intravenously with 5 × 10^3^ CFU *Lm*, and survival was monitored. Mice: (R) WT=5, *Asc*^ΔLysM^*Casp8*^+/−^*Mlkl*^−/−^=5, *Asc*^ΔLysM^*Casp8*^−/−^*Mlkl*^−/−^=8. (S) Graphical summary. Protection against *Fp*^CGD^ is determined not by pyroptosis, apoptosis, or necroptosis themselves, but by antagonistic cytokine signaling embedded within these pathways. RIPK3-dependent type I interferons suppress neutrophil-mediated bacterial clearance and drive susceptibility, whereas caspase-1-dependent IL-1β counteracts this response to restore host resistance. Data and statistical analyses are representative of at least two experiments (A–Q), or pooled from two experiments (R). ns, *p* > 0.05; \*\**p* < 0.01; \*\*\**p* < 0.001; \*\*\*\**p* < 0.0001; two-way ANOVA (A–D, F–I, L, M, O, and Q), one-way ANOVA (N and P), and survival curve analysis (R).

## References

1. Galluzzi, L., Vitale, I., Aaronson, S.A., Abrams, J.M., Adam, D., Agostinis, P., Alnemri, E.S., Altucci, L., Amelio, I., Andrews, D.W., et al. (2018). Molecular mechanisms of cell death: recommendations of the Nomenclature Committee on Cell Death 2018. Cell Death Differ 25, 486–541. 10.1038/s41418-017-0012-4.

2. Bedoui, S., Herold, M.J., and Strasser, A. (2020). Emerging connectivity of programmed cell death pathways and its physiological implications. Nat Rev Mol Cell Bio 21, 678–695. 10.1038/s41580-020-0270-8.

3. Nozaki, K., Li, L., and Miao, E.A. (2022). Innate Sensors Trigger Regulated Cell Death to Combat Intracellular Infection. Annu Rev Immunol 40, 469–498. 10.1146/annurev-immunol-101320-011235.

4. Li, L., Dickinson, M.S., Coers, J., and Miao, E.A. (2023). Pyroptosis in defense against intracellular bacteria. Semin. Immunol. 69, 101805. 10.1016/j.smim.2023.101805.

5. Martinon, F., Burns, K., and Tschopp, J. (2002). The Inflammasome. Mol Cell 10, 417–426. 10.1016/s1097-2765(02)00599-3.

6. Shi, J., Zhao, Y., Wang, K., Shi, X., Wang, Y., Huang, H., Zhuang, Y., Cai, T., Wang, F., and Shao, F. (2015). Cleavage of GSDMD by inflammatory caspases determines pyroptotic cell death. Nature 526, 660–665. 10.1038/nature15514.

7. Thornberry, N.A., and Lazebnik, Y. (1998). Caspases: Enemies Within. Science 281, 1312– 1316. 10.1126/science.281.5381.1312.

8. Cho, Y., Challa, S., Moquin, D., Genga, R., Ray, T.D., Guildford, M., and Chan, F.K.-M. (2009). Phosphorylation-Driven Assembly of the RIP1-RIP3 Complex Regulates Programmed Necrosis and Virus-Induced Inflammation. Cell 137, 1112–1123. 10.1016/j.cell.2009.05.037.

9. Sun, L., Wang, H., Wang, Z., He, S., Chen, S., Liao, D., Wang, L., Yan, J., Liu, W., Lei, X., et al. (2012). Mixed Lineage Kinase Domain-like Protein Mediates Necrosis Signaling Downstream of RIP3 Kinase. Cell 148, 213–227. 10.1016/j.cell.2011.11.031.

10. Lacey, C.A., and Miao, E.A. (2019). Programmed Cell Death in the Evolutionary Race against Bacterial Virulence Factors. Csh Perspect Biol 12, a036459. 10.1101/cshperspect.a036459.

11. Doerflinger, M., Deng, Y., Whitney, P., Salvamoser, R., Engel, S., Kueh, A.J., Tai, L., Bachem, A., Gressier, E., Geoghegan, N.D., et al. (2020). Flexible Usage and Interconnectivity of Diverse Cell Death Pathways Protect against Intracellular Infection. Immunity 53, 533–547.e7. 10.1016/j.immuni.2020.07.004.

12. Roncaioli, J.L., Babirye, J.P., Chavez, R.A., Liu, F.L., Turcotte, E.A., Lee, A.Y., Lesser, C.F., and Vance, R.E. (2023). A hierarchy of cell death pathways confers layered resistance to shigellosis in mice. eLife 12, e83639. 10.7554/elife.83639.

13. Jorgensen, I., Rayamajhi, M., and Miao, E.A. (2017). Programmed cell death as a defence against infection. Nature Publishing Group 17, 151–164. 10.1038/nri.2016.147.

14. Broz, P., and Dixit, V.M. (2016). Inflammasomes: mechanism of assembly, regulation and signalling. Nat Rev Immunol 16, 407–420. 10.1038/nri.2016.58.

15. Maltez, V.I., and Miao, E.A. (2016). Reassessing the Evolutionary Importance of Inflammasomes. The Journal of Immunology 196, 956–962. 10.4049/jimmunol.1502060.

16. Kayagaki, N., Wong, M.T., Stowe, I.B., Ramani, S.R., Gonzalez, L.C., Akashi-Takamura, S., Miyake, K., Zhang, J., Lee, W.P., Muszynski, A., et al. (2013). Noncanonical Inflammasome Activation by Intracellular LPS Independent of TLR4. Science 341, 1246–1249. 10.1126/science.1240248.

17. Hagar, J.A., Powell, D.A., Aachoui, Y., Ernst, R.K., and Miao, E.A. (2013). Cytoplasmic LPS Activates Caspase-11: Implications in TLR4-Independent Endotoxic Shock. Science 341, 1250–1253. 10.1126/science.1240988.

18. Maltez, V.I., Tubbs, A.L., Cook, K.D., Aachoui, Y., Falcone, E.L., Holland, S.M., Whitmire, J.K., and Miao, E.A. (2015). Inflammasomes Coordinate Pyroptosis and Natural Killer Cell Cytotoxicity to Clear Infection by a Ubiquitous Environmental Bacterium. Immunity 43, 987– 997. 10.1016/j.immuni.2015.10.010.

19. Nozaki, K., Maltez, V.I., Rayamajhi, M., Tubbs, A.L., Mitchell, J.E., Lacey, C.A., Harvest, C.K., Li, L., Nash, W.T., Larson, H.N., et al. (2022). Caspase-7 activates ASM to repair gasdermin and perforin pores. Nature 606, 960–967. 10.1038/s41586-022-04825-8.

20. Nozaki, K., and Miao, E.A. (2023). Bucket lists must be completed during cell death. Trends Cell Biol. 10.1016/j.tcb.2023.02.008.

21. Harvest, C.K., Abele, T.J., Yu, C., Beatty, C.J., Amason, M.E., Billman, Z.P., DePrizio, M.A., Souza, F.W., Lacey, C.A., Maltez, V.I., et al. (2023). An innate granuloma eradicates an environmental pathogen using Gsdmd and Nos2. Nat. Commun. 14, 6686. 10.1038/s41467-023-42218-1.

22. Amason, M.E., Beatty, C.J., Harvest, C.K., Saban, D.R., and Miao, E.A. (2024). Chemokine expression profile of an innate granuloma. eLife 13. 10.7554/elife.96425.

23. Winkelstein, J.A., Marino, M.C., Johnston, R.B., Boyle, J., Curnutte, J., Gallin, J.I., Malech, H.L., Holland, S.M., Ochs, H., Quie, P., et al. (2000). Chronic Granulomatous Disease: Report on a National Registry of 368 Patients. Medicine 79, 155–169. 10.1097/00005792-200005000-00003.

24. Holland, S.M. (2010). Chronic Granulomatous Disease. Clin. Rev. Allergy Immunol. 38, 3– 10. 10.1007/s12016-009-8136-z.

25. Nano, F.E., Zhang, N., Cowley, S.C., Klose, K.E., Cheung, K.K.M., Roberts, M.J., Ludu, J.S., Letendre, G.W., Meierovics, A.I., Stephens, G., et al. (2004). A Francisella tularensis Pathogenicity Island Required for Intramacrophage Growth. J. Bacteriol. 186, 6430–6436. 10.1128/jb.186.19.6430-6436.2004.

26. Dennis, D.T., Inglesby, T.V., Henderson, D.A., Bartlett, J.G., Ascher, M.S., Eitzen, E., Fine, A.D., Friedlander, A.M., Hauer, J., Layton, M., et al. (2001). Tularemia as a Biological Weapon: Medical and Public Health Management. Jama 285, 2763–2773. 10.1001/jama.285.21.2763.

27. Wenger, J.D., Hollis, D.G., Weaver, R.E., Baker, C.N., Brown, G.R., Brenner, D.J., and Broome, C.V. (1989). Infection Caused by Francisella philomiragia (Formerly Yersinia philomiragia): A Newly Recognized Human Pathogen. Ann Intern Med 110, 888. 10.7326/0003-4819-110-11-888.

28. Friis-Møller, A., Lemming, L.E., Valerius, N.H., and Bruun, B. (2004). Problems in Identification of Francisella philomiragia Associated with Fatal Bacteremia in a Patient with Chronic Granulomatous Disease. J Clin Microbiol 42, 1840–1842. 10.1128/jcm.42.4.1840-1842.2004.

29. Timothy, M., and H, S. (2005). Francisella philomiragia Adenitis and Pulmonary Nodules in a Child with Chronic Granulomatous Disease. Can J Infect Dis Medical Microbiol 16, 245– 248. 10.1155/2005/486417.

30. Amason, M.E., Li, L., Harvest, C.K., Lacey, C.A., and Miao, E.A. (2024). Validation of the Intermolecular Disulfide Bond in Caspase-2. Biology 13, 49. 10.3390/biology13010049.

31. Kocks, C., Gouin, E., Tabouret, M., Berche, P., Ohayon, H., and Cossart, P. (1992). L. monocytogenes-induced actin assembly requires the actA gene product, a surface protein. Cell 68, 521–531. 10.1016/0092-8674(92)90188-i.

32. Jorgensen, I., Zhang, Y., Krantz, B.A., and Miao, E.A. (2016). Pyroptosis triggers pore-induced intracellular traps (PITs) that capture bacteria and lead to their clearance by efferocytosis. The Journal of Experimental Medicine 213, 2113–2128. 10.1084/jem.20151613.

33. Fernandes-Alnemri, T., Yu, J.-W., Datta, P., Wu, J., and Alnemri, E.S. (2009). AIM2 activates the inflammasome and cell death in response to cytoplasmic DNA. Nature 458, 509–513. 10.1038/nature07710.

34. Fernandes-Alnemri, T., Yu, J.-W., Juliana, C., Solorzano, L., Kang, S., Wu, J., Datta, P., McCormick, M., Huang, L., McDermott, E., et al. (2010). The AIM2 inflammasome is critical for innate immunity to Francisella tularensis. Nat Immunol 11, 385–393. 10.1038/ni.1859.

35. Jones, J.W., Kayagaki, N., Broz, P., Henry, T., Newton, K., O’Rourke, K., Chan, S., Dong, J., Qu, Y., Roose-Girma, M., et al. (2010). Absent in melanoma 2 is required for innate immune recognition of Francisella tularensis. Proc National Acad Sci 107, 9771–9776. 10.1073/pnas.1003738107.

36. Rathinam, V.A.K., Jiang, Z., Waggoner, S.N., Sharma, S., Cole, L.E., Waggoner, L., Vanaja, S.K., Monks, B.G., Ganesan, S., Latz, E., et al. (2010). The AIM2 inflammasome is essential for host defense against cytosolic bacteria and DNA viruses. Nat Immunol 11, 395–402. 10.1038/ni.1864.

37. Muñoz-Planillo, R., Kuffa, P., Martínez-Colón, G., Smith, B.L., Rajendiran, T.M., and Núñez, G. (2013). K+ Efflux Is the Common Trigger of NLRP3 Inflammasome Activation by Bacterial Toxins and Particulate Matter. Immunity 38, 1142–1153. 10.1016/j.immuni.2013.05.016.

38. Swanson, K.V., Deng, M., and Ting, J.P.-Y. (2019). The NLRP3 inflammasome: molecular activation and regulation to therapeutics. Nat. Rev. Immunol. 19, 477–489. 10.1038/s41577-019-0165-0.

39. Tummers, B., Mari, L., Guy, C.S., Heckmann, B.L., Rodriguez, D.A., Rühl, S., Moretti, J., Crawford, J.C., Fitzgerald, P., Kanneganti, T.-D., et al. (2020). Caspase-8-Dependent Inflammatory Responses Are Controlled by Its Adaptor, FADD, and Necroptosis. Immunity 52, 994–1006.e8. 10.1016/j.immuni.2020.04.010.

40. Pierini, R., Juruj, C., Perret, M., Jones, C.L., Mangeot, P., Weiss, D.S., and Henry, T. (2012). AIM2/ASC triggers caspase-8-dependent apoptosis in Francisella-infected caspase-1-deficient macrophages. Cell Death Differ 19, 1709–1721. 10.1038/cdd.2012.51.

41. Sagulenko, V., Thygesen, S.J., Sester, D.P., Idris, A., Cridland, J.A., Vajjhala, P.R., Roberts, T.L., Schroder, K., Vince, J.E., Hill, J.M., et al. (2013). AIM2 and NLRP3 inflammasomes activate both apoptotic and pyroptotic death pathways via ASC. Cell Death Differ 20, 1149– 1160. 10.1038/cdd.2013.37.

42. Alvarez-Diaz, S., Dillon, C.P., Lalaoui, N., Tanzer, M.C., Rodriguez, D.A., Lin, A., Lebois, M., Hakem, R., Josefsson, E.C., O’Reilly, L.A., et al. (2016). The Pseudokinase MLKL and the Kinase RIPK3 Have Distinct Roles in Autoimmune Disease Caused by Loss of Death-Receptor-Induced Apoptosis. Immunity 45, 513–526. 10.1016/j.immuni.2016.07.016.

43. Daniels, B.P., Snyder, A.G., Olsen, T.M., Orozco, S., Oguin, T.H., Tait, S.W.G., Martinez, J., Gale, M., Loo, Y.-M., and Oberst, A. (2017). RIPK3 Restricts Viral Pathogenesis via Cell Death-Independent Neuroinflammation. Cell 169, 301–313.e11. 10.1016/j.cell.2017.03.011.

44. Daniels, B.P., Kofman, S.B., Smith, J.R., Norris, G.T., Snyder, A.G., Kolb, J.P., Gao, X., Locasale, J.W., Martinez, J., Gale, M., et al. (2019). The Nucleotide Sensor ZBP1 and Kinase RIPK3 Induce the Enzyme IRG1 to Promote an Antiviral Metabolic State in Neurons. Immunity 50, 64–76.e4. 10.1016/j.immuni.2018.11.017.

45. Kofman, S.B., Chu, L.H., Ames, J.M., Chavarria, S.D., Lichauco, K., Daniels, B.P., and Oberst, A. (2025). RIPK3 coordinates RHIM domain–dependent antiviral inflammatory transcription in neurons. Sci. Signal. 18, eado9745. 10.1126/scisignal.ado9745.

46. Oberst, A., Dillon, C.P., Weinlich, R., McCormick, L.L., Fitzgerald, P., Pop, C., Hakem, R., Salvesen, G.S., and Green, D.R. (2011). Catalytic activity of the caspase-8–FLIPL complex inhibits RIPK3-dependent necrosis. Nature 471, 363–367. 10.1038/nature09852.

47. Brown, D.E., Libby, S.J., Moreland, S.M., McCoy, M.W., Brabb, T., Stepanek, A., Fang, F.C., and Detweiler, C.S. (2013). Salmonella enterica Causes More Severe Inflammatory Disease in C57/BL6 Nramp1 G169Mice Than Sv129S6 Mice. Veterinary Pathology 50, 867–876. 10.1177/0300985813478213.

48. Dougan, G., John, V., Palmer, S., and Mastroeni, P. (2011). Immunity to salmonellosis. Immunol. Rev. 240, 196–210. 10.1111/j.1600-065x.2010.00999.x.

49. Robbins, J.R., Barth, A.I., Marquis, H., Hostos, E.L. de, Nelson, W.J., and Theriot, J.A. (1999). Listeria monocytogenes Exploits Normal Host Cell Processes to Spread from Cell to Cell✪. J. Cell Biol. 146, 1333–1350. 10.1083/jcb.146.6.1333.

50. Sims, J.E., and Smith, D.E. (2010). The IL-1 family: regulators of immunity. Nat Rev Immunol 10, 89–102. 10.1038/nri2691.

51. Boxx, G.M., and Cheng, G. (2016). The Roles of Type I Interferon in Bacterial Infection. Cell Host Microbe 19, 760–769. 10.1016/j.chom.2016.05.016.

52. Reizis, B. (2019). Plasmacytoid Dendritic Cells: Development, Regulation, and Function. Immunity 50, 37–50. 10.1016/j.immuni.2018.12.027.

53. Guo, H., Kaiser, W.J., and Mocarski, E.S. (2015). Manipulation of apoptosis and necroptosis signaling by herpesviruses. Méd. Microbiol. Immunol. 204, 439–448. 10.1007/s00430-015-0410-5.

54. McNab, F., Mayer-Barber, K., Sher, A., Wack, A., and O’Garra, A. (2015). Type I interferons in infectious disease. Nat Rev Immunol 15, 87–103. 10.1038/nri3787.

55. Nguyen, G.T., Green, E.R., and Mecsas, J. (2017). Neutrophils to the ROScue: Mechanisms of NADPH Oxidase Activation and Bacterial Resistance. Front. Cell. Infect. Microbiol. 7, 373. 10.3389/fcimb.2017.00373.

56. Canton, M., Sánchez-Rodríguez, R., Spera, I., Venegas, F.C., Favia, M., Viola, A., and Castegna, A. (2021). Reactive Oxygen Species in Macrophages: Sources and Targets. Front. Immunol. 12, 734229. 10.3389/fimmu.2021.734229.

57. Kurihara, T., Warr, G., Loy, J., and Bravo, R. (1997). Defects in Macrophage Recruitment and Host Defense in Mice Lacking the CCR2 Chemokine Receptor. J. Exp. Med. 186, 1757– 1762. 10.1084/jem.186.10.1757.

58. Gurung, P., Anand, P.K., Malireddi, R.K.S., Walle, L.V., Opdenbosch, N.V., Dillon, C.P., Weinlich, R., Green, D.R., Lamkanfi, M., and Kanneganti, T.-D. (2014). FADD and Caspase-8 Mediate Priming and Activation of the Canonical and Noncanonical Nlrp3 Inflammasomes. J. Immunol. 192, 1835–1846. 10.4049/jimmunol.1302839.

59. Antonopoulos, C., Russo, H.M., Sanadi, C.E., Martin, B.N., Li, X., Kaiser, W.J., Mocarski, E.S., and Dubyak, G.R. (2015). Caspase-8 as an Effector and Regulator of NLRP3 Inflammasome Signaling*. J Biol Chem 290, 20167–20184. 10.1074/jbc.m115.652321.

60. Case, C.L., Kohler, L.J., Lima, J.B., Strowig, T., Zoete, M.R. de, Flavell, R.A., Zamboni, D.S., and Roy, C.R. (2013). Caspase-11 stimulates rapid flagellin-independent pyroptosis in response to Legionella pneumophila. Proc National Acad Sci 110, 1851–1856. 10.1073/pnas.1211521110.

61. Martin, B.N., Wang, C., Zhang, C., Kang, Z., Gulen, M.F., Zepp, J.A., Zhao, J., Bian, G., Do, J., Min, B., et al. (2016). T cell–intrinsic ASC critically promotes TH17-mediated experimental autoimmune encephalomyelitis. Nat. Immunol. 17, 583–592. 10.1038/ni.3389.

62. Guarda, G., Braun, M., Staehli, F., Tardivel, A., Mattmann, C., Förster, I., Farlik, M., Decker, T., Du Pasquier, R.A., Romero, P., et al. (2011). Type I Interferon Inhibits Interleukin-1 Production and Inflammasome Activation. Immunity 34, 213–223. 10.1016/j.immuni.2011.02.006.

63. Garlanda, C., Dinarello, C.A., and Mantovani, A. (2013). The Interleukin-1 Family: Back to the Future. Immunity 39, 1003–1018. 10.1016/j.immuni.2013.11.010.

64. Moriwaki, K., and Chan, F.K.-M. (2013). RIP3: a molecular switch for necrosis and inflammation. Genes Dev. 27, 1640–1649. 10.1101/gad.223321.113.

65. Rodriguez, D.A., Quarato, G., Liedmann, S., Tummers, B., Zhang, T., Guy, C., Crawford, J.C., Palacios, G., Pelletier, S., Kalkavan, H., et al. (2022). Caspase-8 and FADD prevent spontaneous ZBP1 expression and necroptosis. Proc National Acad Sci 119, e2207240119. 10.1073/pnas.2207240119.

66. Kelepouras, K., Saggau, J., Bonasera, D., Kiefer, C., Locci, F., Rakhsh-Khorshid, H., Grauvogel, L., Varanda, A.B., Peifer, M., Loricchio, E., et al. (2025). STING induces ZBP1-mediated necroptosis independently of TNFR1 and FADD. Nature 647, 735–746. 10.1038/s41586-025-09536-4.

67. Feng, S., Yang, Y., Mei, Y., Ma, L., Zhu, D., Hoti, N., Castanares, M., and Wu, M. (2007). Cleavage of RIP3 inactivates its caspase-independent apoptosis pathway by removal of kinase domain. Cell. Signal. 19, 2056–2067. 10.1016/j.cellsig.2007.05.016.

68. Abram, C.L., Roberge, G.L., Hu, Y., and Lowell, C.A. (2014). Comparative analysis of the efficiency and specificity of myeloid-Cre deleting strains using ROSA-EYFP reporter mice. J Immunol Methods 408, 89–100. 10.1016/j.jim.2014.05.009.

69. Jones, J.D.G., and Dangl, J.L. (2006). The plant immune system. Nature 444, 323–329. 10.1038/nature05286.

70. Thome, M., Schneider, P., Hofmann, K., Fickenscher, H., Meinl, E., Neipel, F., Mattmann, C., Burns, K., Bodmer, J.-L., Schröter, M., et al. (1997). Viral FLICE-inhibitory proteins (FLIPs) prevent apoptosis induced by death receptors. Nature 386, 517–521. 10.1038/386517a0.

71. Upton, J.W., Kaiser, W.J., and Mocarski, E.S. (2010). Virus Inhibition of RIP3-Dependent Necrosis. Cell Host Microbe 7, 302–313. 10.1016/j.chom.2010.03.006.

72. Dondelinger, Y., Hulpiau, P., Saeys, Y., Bertrand, M.J.M., and Vandenabeele, P. (2016). An evolutionary perspective on the necroptotic pathway. Trends Cell Biol. 26, 721–732. 10.1016/j.tcb.2016.06.004.

73. Auerbuch, V., Brockstedt, D.G., Meyer-Morse, N., O’Riordan, M., and Portnoy, D.A. (2004). Mice Lacking the Type I Interferon Receptor Are Resistant to Listeria monocytogenes. J. Exp. Med. 200, 527–533. 10.1084/jem.20040976.

74. Mayer-Barber, K.D., Andrade, B.B., Oland, S.D., Amaral, E.P., Barber, D.L., Gonzales, J., Derrick, S.C., Shi, R., Kumar, N.P., Wei, W., et al. (2014). Host-directed therapy of tuberculosis based on interleukin-1 and type I interferon crosstalk. Nature 511, 99–103. 10.1038/nature13489.

75. Wang, Y., Deng, Y., Chen, J., Hahn, Q., Umbaugh, D.S., Zhang, Z., Zhang, Y., Rowe, S.E., Li, L., Herring, L.E., et al. (2025). cGAS Inhibits ALDH2 to Suppress Lipid Droplet Function and Regulate MASLD Progression. Adv. Sci. 12, e08576. 10.1002/advs.202508576.

76. Wells, A.I., Grimes, K.A., and Coyne, C.B. (2022). Enterovirus Replication and Dissemination Are Differentially Controlled by Type I and III Interferons in the Gastrointestinal Tract. mBio 13, e00443–22. 10.1128/mbio.00443-22.

77. Rowe, S.E., Wagner, N.J., Li, L., Beam, J.E., Wilkinson, A.D., Radlinski, L.C., Zhang, Q., Miao, E.A., and Conlon, B.P. (2020). Reactive oxygen species induce antibiotic tolerance during systemic Staphylococcus aureus infection. Nat Microbiol 5, 282–290. 10.1038/s41564-019-0627-y.

78. Mariathasan, S., Weiss, D.S., Newton, K., McBride, J., O’Rourke, K., Roose-Girma, M., Lee, W.P., Weinrauch, Y., Monack, D.M., and Dixit, V.M. (2006). Cryopyrin activates the inflammasome in response to toxins and ATP. Nature 440, 228–232. 10.1038/nature04515.

79. Oh, C., Li, L., Verma, A., Reuven, A.D., Miao, E.A., Bliska, J.B., and Aachoui, Y. (2022). Neutrophil inflammasomes sense the subcellular delivery route of translocated bacterial effectors and toxins. Cell Reports 41, 111688. 10.1016/j.celrep.2022.111688.

80. Aachoui, Y., Kajiwara, Y., Leaf, I.A., Mao, D., Ting, J.P.-Y., Coers, J., Aderem, A., Buxbaum, J.D., and Miao, E.A. (2015). Canonical Inflammasomes Drive IFN-γ to Prime Caspase-11 in Defense against a Cytosol-Invasive Bacterium. Cell Host & Microbe 18, 320–332. 10.1016/j.chom.2015.07.016.

81. Aachoui, Y., Leaf, I.A., Hagar, J.A., Fontana, M.F., Campos, C.G., Zak, D.E., Tan, M.H., Cotter, P.A., Vance, R.E., Aderem, A., et al. (2013). Caspase-11 Protects Against Bacteria That Escape the Vacuole. Science 339, 1230751–1230978. 10.1126/science.1230751.

82. Abele, T.J., Billman, Z.P., Li, L., Harvest, C.K., Bryan, A.K., Magalski, G.R., Lopez, J.P., Larson, H.N., Yin, X.-M., and Miao, E.A. (2023). Apoptotic signaling clears engineered Salmonella in an organ-specific manner. 10.1101/2023.05.06.539681.

83. Murphy, J.M., Czabotar, P.E., Hildebrand, J.M., Lucet, I.S., Zhang, J.-G., Alvarez-Diaz, S., Lewis, R., Lalaoui, N., Metcalf, D., Webb, A.I., et al. (2013). The Pseudokinase MLKL Mediates Necroptosis via a Molecular Switch Mechanism. Immunity 39, 443–453. 10.1016/j.immuni.2013.06.018.

84. Newton, K., Sun, X., and Dixit, V.M. (2004). Kinase RIP3 Is Dispensable for Normal NF-κBs, Signaling by the B-Cell and T-Cell Receptors, Tumor Necrosis Factor Receptor 1, and Toll-Like Receptors 2 and 4. Mol Cell Biol *24*, 1464–1469. 10.1128/mcb.24.4.1464-1469.2004.

85. Moriwaki, K., Balaji, S., Bertin, J., Gough, P.J., and Chan, F.K.-M. (2017). Distinct Kinase-Independent Role of RIPK3 in CD11c+ Mononuclear Phagocytes in Cytokine-Induced Tissue Repair. Cell Rep. 18, 2441–2451. 10.1016/j.celrep.2017.02.015.

86. Dillon, C.P., Weinlich, R., Rodriguez, D.A., Cripps, J.G., Quarato, G., Gurung, P., Verbist, K.C., Brewer, T.L., Llambi, F., Gong, Y.-N., et al. (2014). RIPK1 Blocks Early Postnatal Lethality Mediated by Caspase-8 and RIPK3. Cell 157, 1189–1202. 10.1016/j.cell.2014.04.018.

87. Mariathasan, S., Newton, K., Monack, D.M., Vucic, D., French, D.M., Lee, W.P., Roose-Girma, M., Erickson, S., and Dixit, V.M. (2004). Differential activation of the inflammasome by caspase-1 adaptors ASC and Ipaf. Nature 430, 213–218. 10.1038/nature02664.

88. Kuida, K., Lippke, J.A., Ku, G., Harding, M.W., Livingston, D.J., Su, M.S.-S., and Flavell, R.A. (1995). Altered Cytokine Export and Apoptosis in Mice Deficient in Interleukin-1β Converting Enzyme. Science 267, 2000–2003. 10.1126/science.7535475.

89. Li, L., Kovacs, S.B., Jørgensen, I., Larson, H.N., Lazear, H.M., and Miao, E.A. (2022). Role of Caspases and Gasdermin A during HSV-1 Infection in Mice. Viruses 14, 2034. 10.3390/v14092034.

90. Dong, H., Zhao, B., Chen, J., Liu, Z., Li, X., Li, L., and Wen, H. (2022). Mitochondrial calcium uniporter promotes phagocytosis-dependent activation of the NLRP3 inflammasome. Proc National Acad Sci 119, e2123247119. 10.1073/pnas.2123247119.

91. Guo, J. (Sammy), Lakhani, F.K., Vientos, C.G., Li, L., and Miao, E.A. (2026). Salmonella SPI2 evades detection by NAIP/NLRC4 despite utilization of a detectable needle. bioRxiv, 2026.01.05.696446. 10.64898/2026.01.05.696446.

