## Supplementary figures and images for "Cytokines driven by caspase-1 and RIPK3 are antagonistic"

### Figure S1

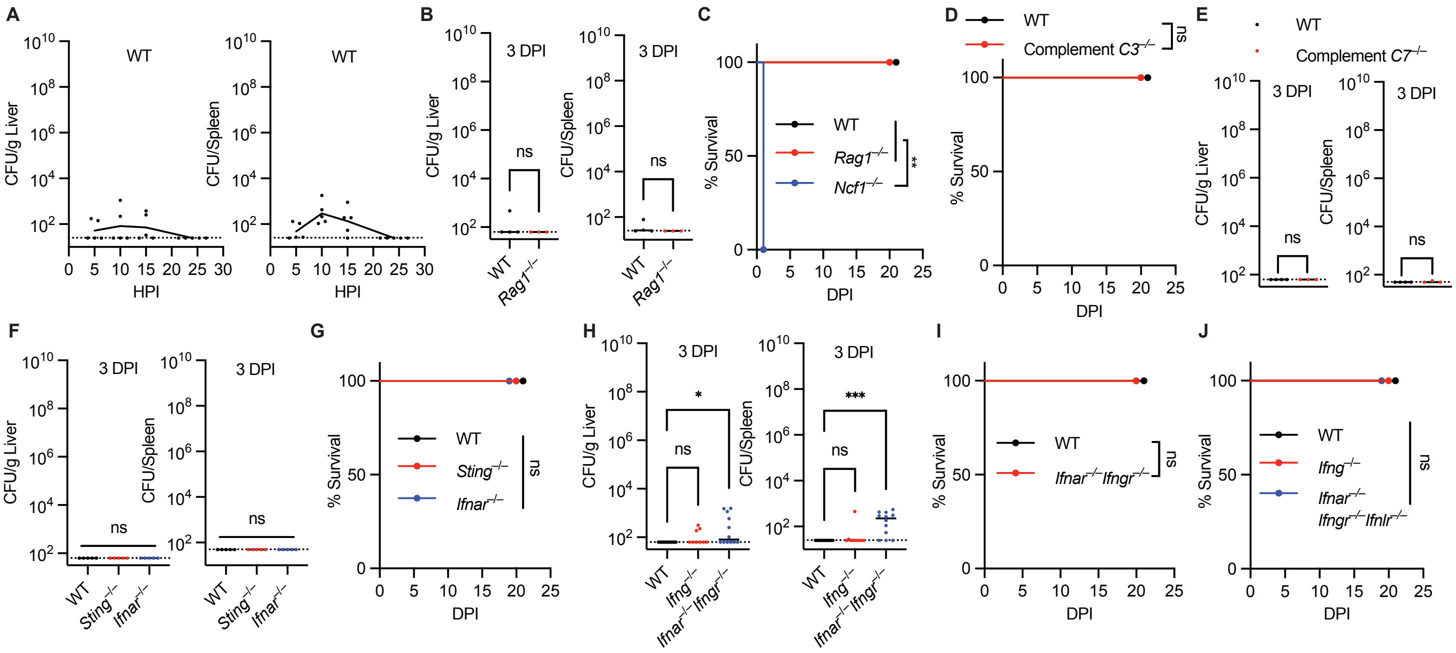

### Figure S2

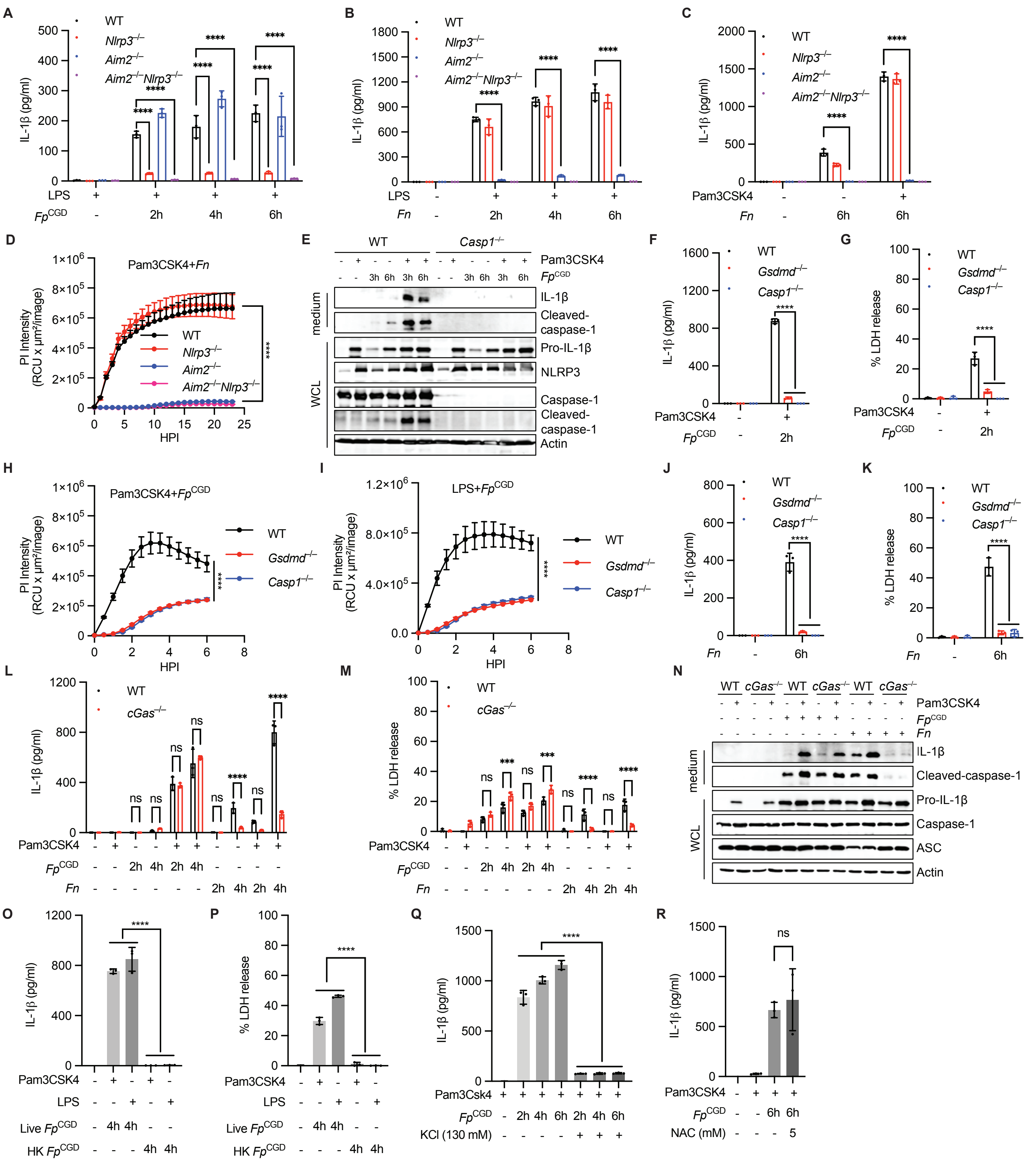

### Figure S3

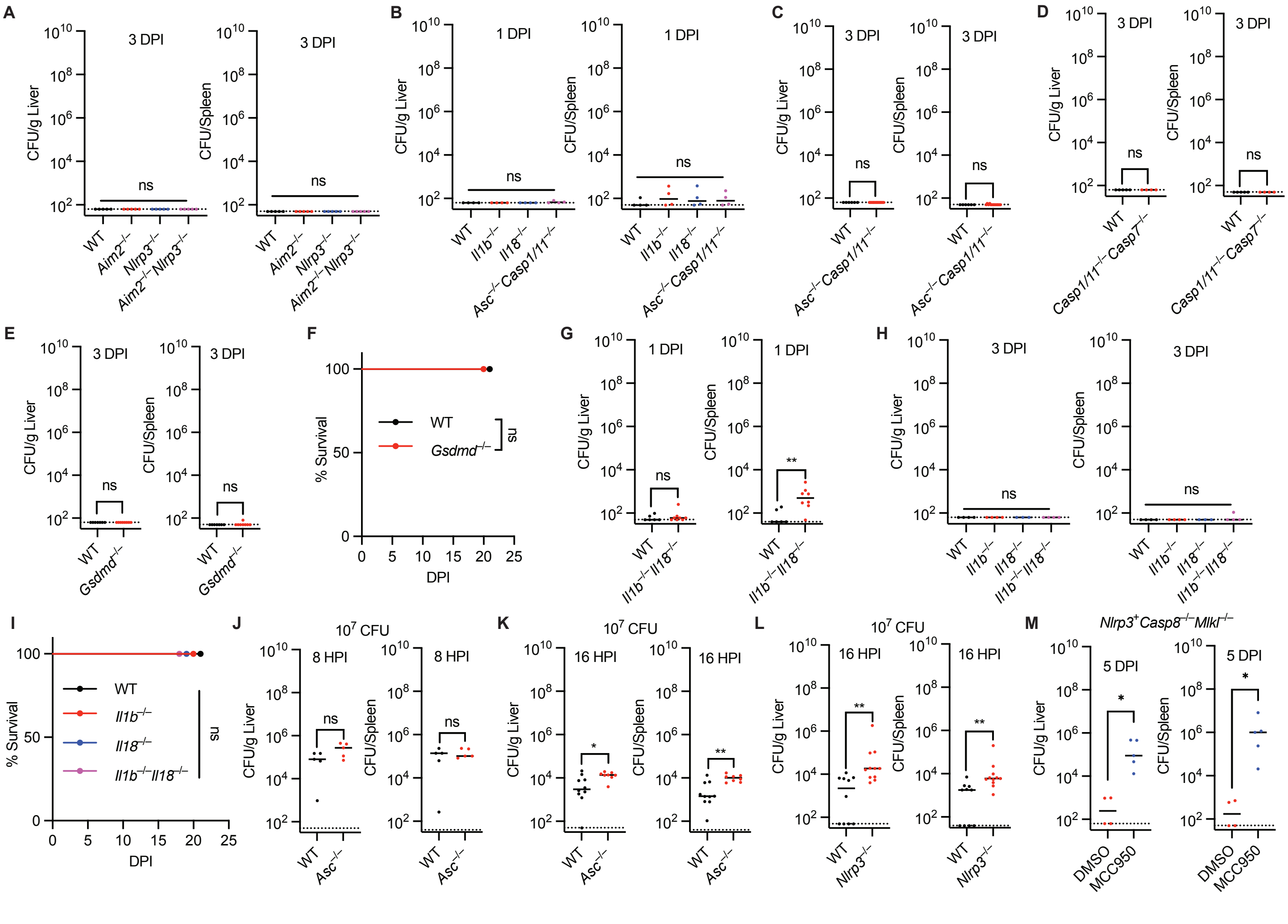

### Figure S4

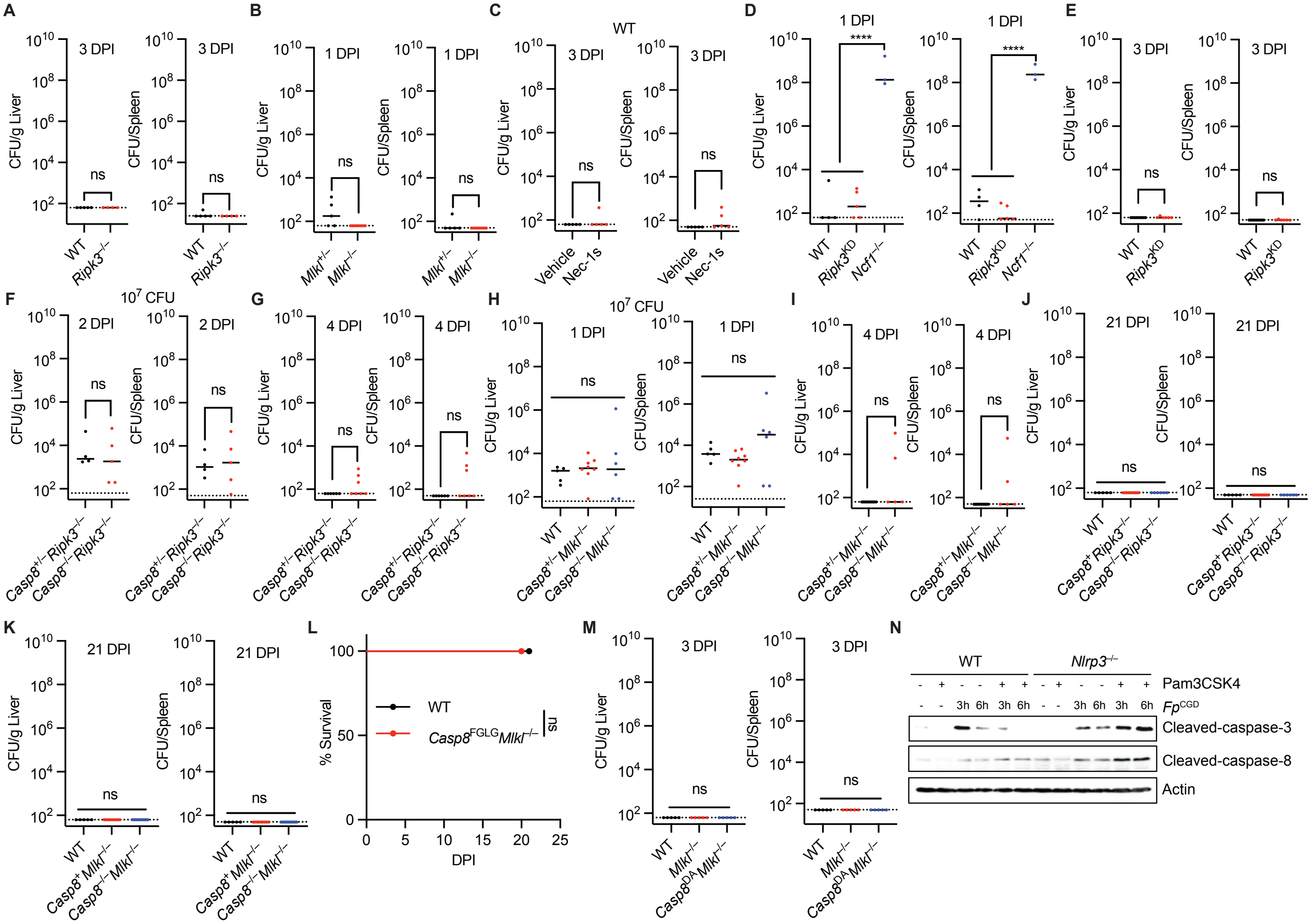

### Figure S5

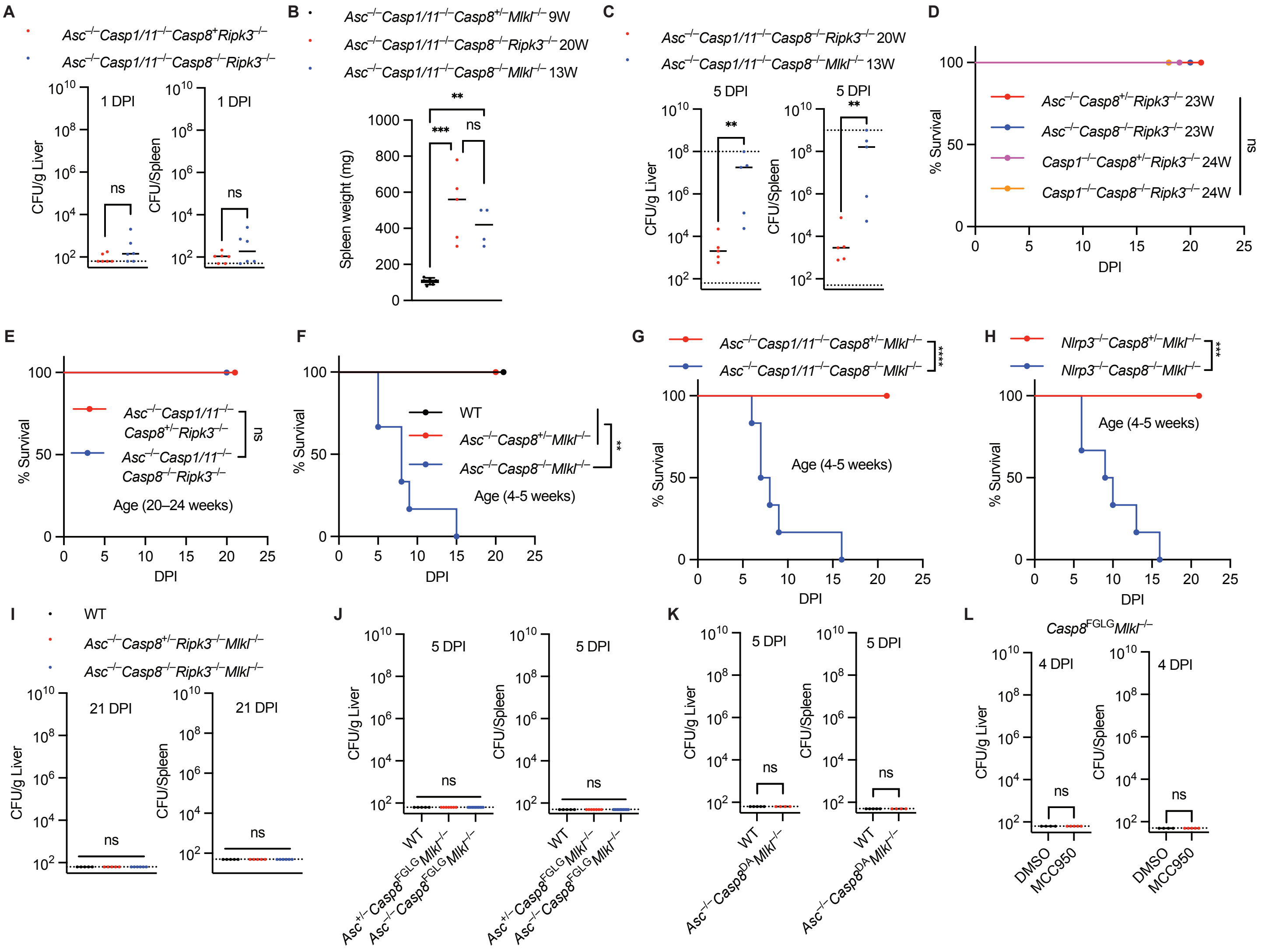

### Figure S6

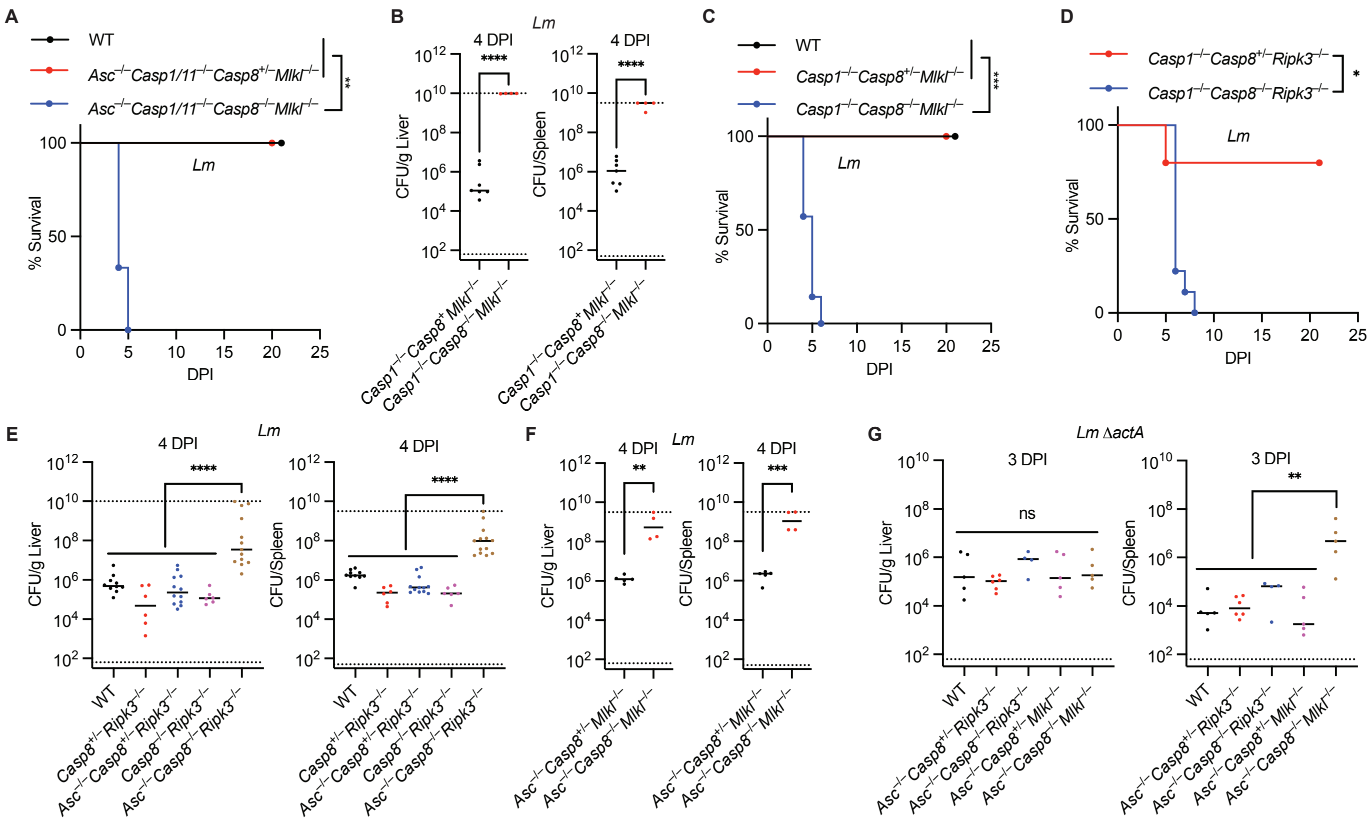

### Figure S7

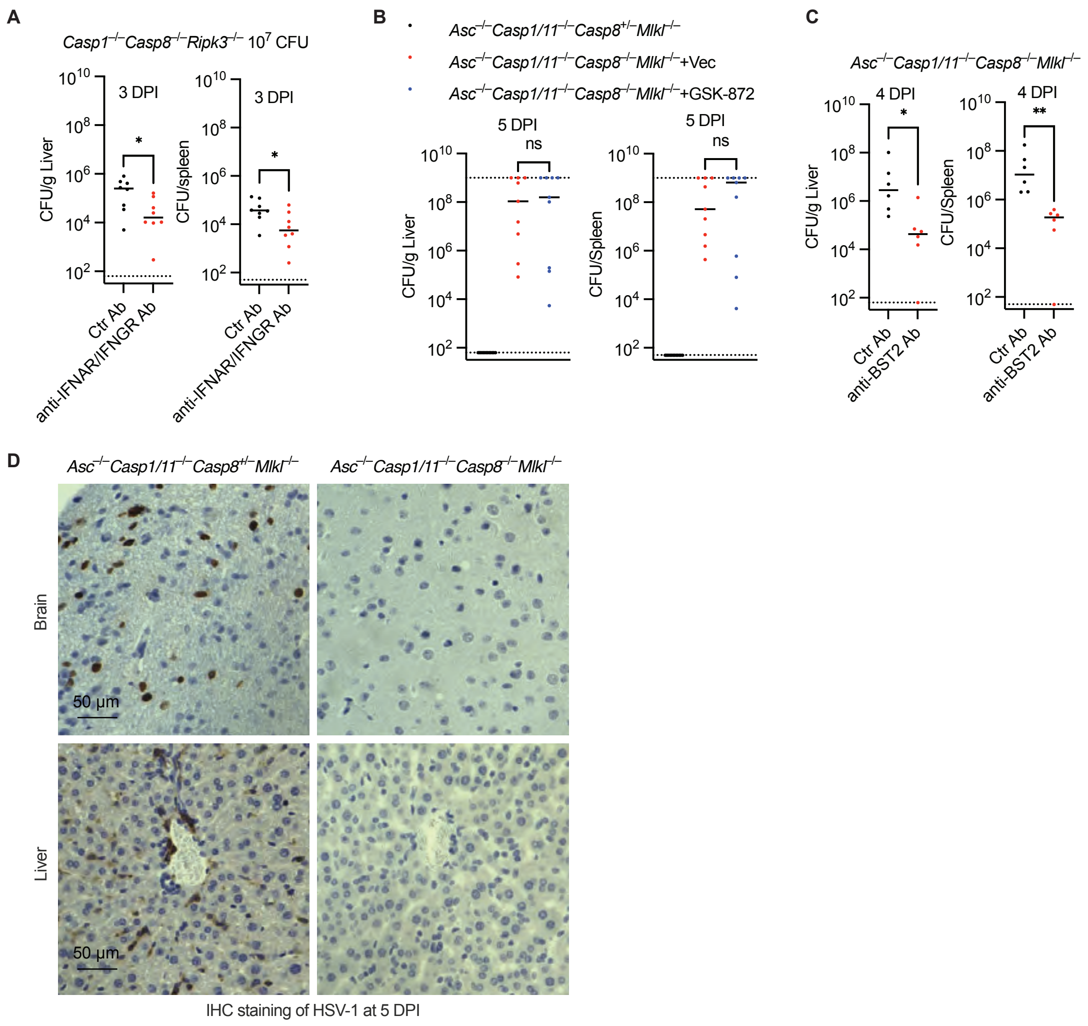

### Figure S8

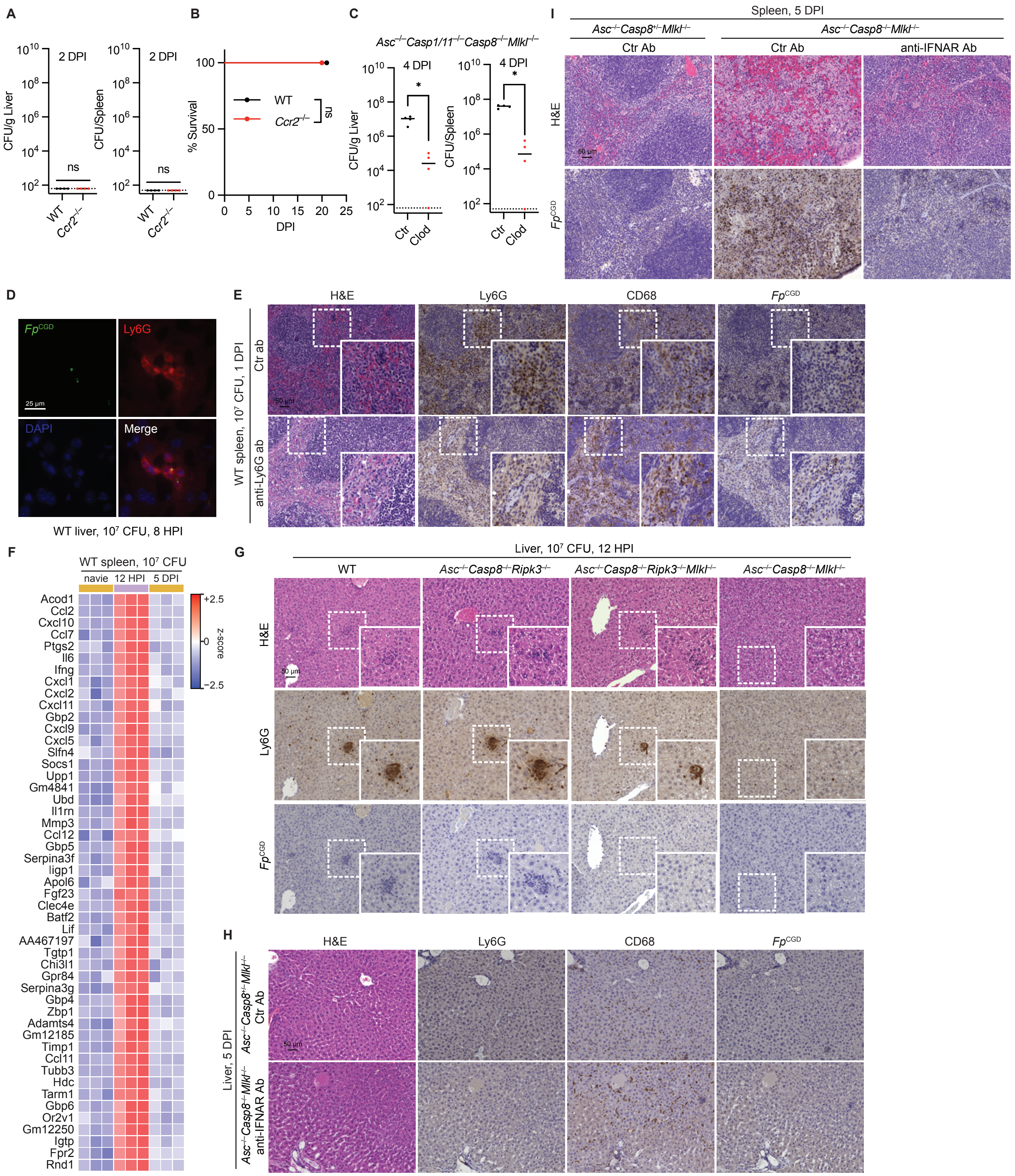

### Figure S9

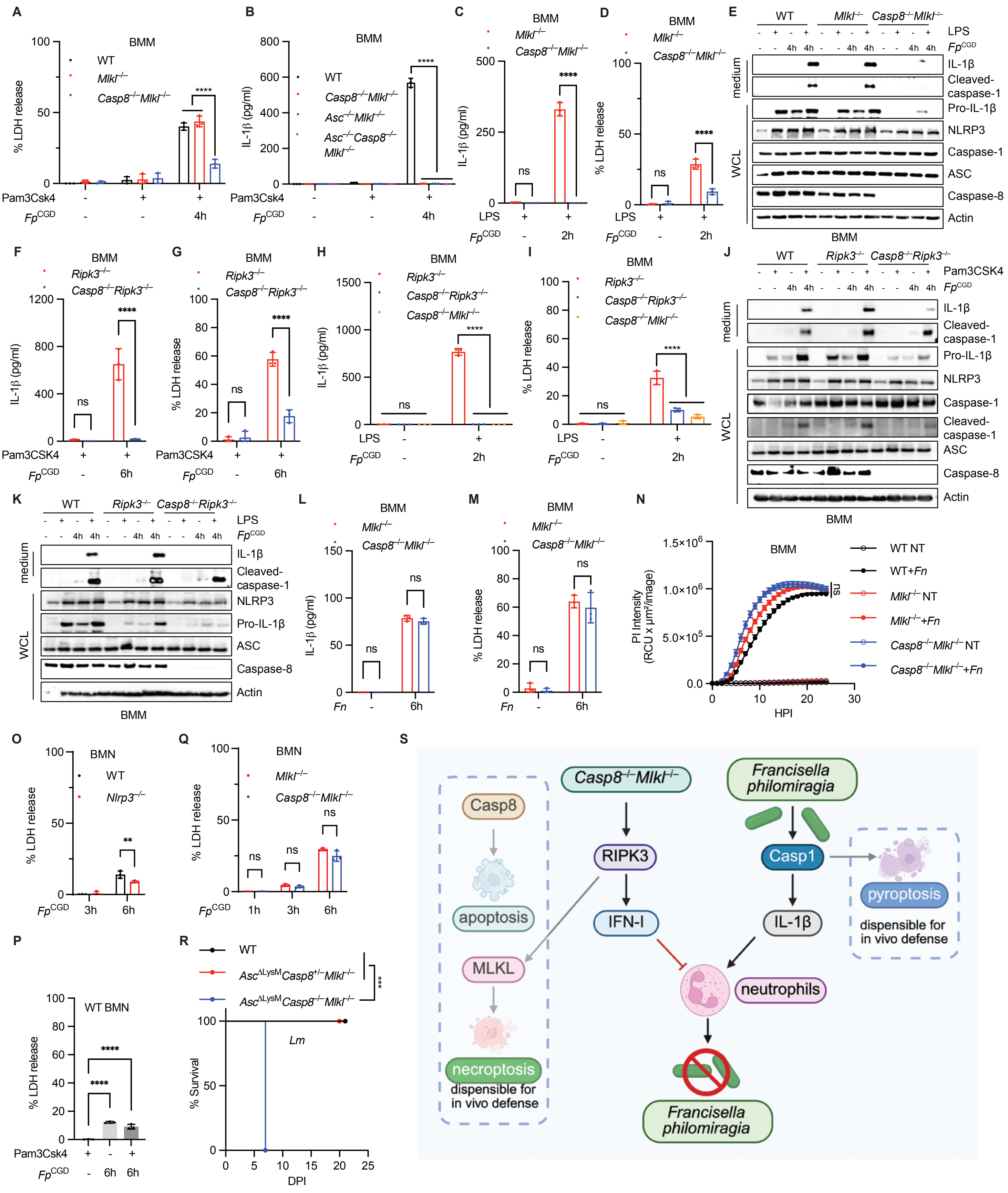
